# Adhesion of *E. coli* bacteria is force-modulated due to fimbriae-mediated surface repulsion and multivalent binding irrespective of surface specificity

**DOI:** 10.1101/2024.05.23.595589

**Authors:** Anders Lundgren, Peter van Oostrum, Jagoba Iturri, Michael Malkoch, José Luis Toca-Herrara, Erik Reimhult

**Affiliations:** Department of Chemistry and Molecular Biology, University of Gothenburg, 405 30 Gothenburg, Sweden; Institute of Colloid and Biointerface Science (ICBS), Department of Bionanosciences (DBNS), BOKU University, 1190 Vienna, Austria; Institute of Biophysics, Department of Bionanosciences (DBNS), BOKU University, 1190 Vienna, Austria; Department of Fibre and Polymer Technology, KTH Royal Institute of Technology, 100 44 Stockholm, Sweden; Centre for Antibiotic Resistance Research (CARe), University of Gothenburg, 41346 Gothenburg, Sweden

## Abstract

*Escherichia coli* bacteria that express type 1 fimbriae migrate along surfaces when pushed by a slow flow but stick more firmly when the flow increases. This and other examples of force-modulated biological binding are often described as due to lectin–glycan catch-bonds. Here we quantitatively track the 3D movements of fimbriated *E. coli* flowing over surfaces nanopatterned with mannose or hydrophobic binding sites. We reveal that flow-modulated surface adhesion and motion are consequences of bacteria adhering via polydisperse, elastic fimbriae, irrespective of binding affinity and specificity. The fimbria-mediated surface repulsion and the flow forces on tethered bacteria establish an equilibrium bacteria-surface separation. The separation controls the number of potential tethers between the bacterium and the surface. Combined with the individual fimbria affinity, this determines the surface avidity and surface motion. This provides a broadly applicable mechanism by which bacteria acquire adaptive surface avidity, responding super-selectively to different flow environments, concentration, and affinity of available binding sites, essential to explaining how fimbriae govern tropism and surface colonization.

## Introduction

Type 1 fimbriae (or type 1 pili) are the most studied chaperon-usher (CU) pili. These are structurally resembling proteinaceous extensions from the bacterial body found on many Gram-negative bacteria, particularly in *Enterobacteriaceae*, e.g., *E. coli*^1,2^. CU pili are important for pathogenesis, e.g., biofilm formation, and as determinants of bacterial tropism^3–11^. An *E. coli* bacterium typically expresses hundreds of type 1 fimbriae with lengths varying between a few hundred nanometers to several micrometers^12,13^. Each fimbria consists of thousands of copies of the FimA protein assembled into a 7 nm-wide helically-coiled rod that presents, distally, the mannose-binding lectin FimH^14,15^. The type 1-fimbriated *E. coli* can form weak transient surface bonds that give rise to the so-called “stick-and-roll motion” in a slow flow, while the bonds drastically strengthen when the flow accelerates. The force-modulated binding is considered a hallmark of the specific catch-bond that forms between FimH and mannosylated proteins exposed on tissue surfaces^16–19^. This bond type, in synergy with fimbriae extension due to reversible uncoiling of the fimbrial rod^20–24^, is believed to aid bacteria in staying surface-bound when exposed to strong flow and thereby promote biofilm formation in the urinary tract; *E. coli* is indeed the most common cause of urinary tract infections^25^. Many studies have detailed the FimH–mannose interaction, and anti-adhesion therapies aiming for the competitive displacement of bacteria by molecular mannose or derivates have been suggested as an alternative to classical antibiotic drugs^26–33^. Yet, the exclusive dependence of force-modulated binding on the FimH-mannose bond, or even on catch-bonds in general, is questionable. Similar binding behavior was observed for *E. coli* expressing P-fimbriae^34^ that bind to digalactose *via* slip bonds^35^. Furthermore, the infectivity of *E. coli* in mice depends strongly on the fimbrial shaft properties in studies where this, but not the function of FimH, is altered^36,37^. Contrary to what one would expect, type 1 fimbriae are indispensable for biofilm formation also on abiotic surfaces like catheters and other biomaterials that do not present mannose^38–40^. Together, these observations imply a more general mechanism for how fimbriae contribute to bacterial binding and biofilm formation than explained purely by the specific FimH– mannose interaction.

A critical aspect of the adhesion of fimbriated bacteria is that a multiplicity of fimbriae may partake in binding. The avidity of surface-bound objects increases sharply with the number of tethers because the probability of all bonds simultaneously releasing drops dramatically with increasing tether multivalency^41,42^. Theoretical work and computer simulations show that the flow-modulated binding of fimbriated bacteria^43,44^, blood cells^45–48^, and artificial particles^49,50^ indeed may depend on a force-induced shift in the number of surface tethers. It is challenging to prove this experimentally since it requires nanometer-level 3D-positioning of a moving microscopic object *and* the simultaneous measurement of the flow forces. In addition, the number of surface tethers must be either measured or controlled. Previous works demonstrate the use of reflection interference contrast microscopy for high-speed 3D positioning of lymphocytes that bound transiently to surfaces^51^ and holographic microscopy to reconstruct 3D trajectories of predominately swimming, flagellated bacteria^52–55^. Here we used an adaptation of holographic microscopy to track the movements of every single type-1 fimbriated *E. coli* (without flagella) flowing through a microfluidic channel. The non-binding bacteria’s trajectories were used to estimate the flow force acting on the surface-interacting bacteria. We also nanopatterned the channels with mannose or hydrophobic binding sites to control the number of specific or non-specific fimbrial surface contacts that a bacterium could form. We show that due to the multivalent binding capacity, irrespective of the surface specificity, fimbriae make bacterial binding super-selective *and* force-modulated through a mechanism that relies on the balance between the repulsive and the binding properties of fimbriae. These findings provide a universal explanation how type 1 fimbriae, or other structurally similar pili, can modulate bacterial colonization.

## Results

### Fimbriae lend bacteria both repulsive and adhesive surface interaction irrespective of the surface specificity

Using electron microscopy (EM) and atomic force microscopy (AFM), we characterized the appearance and surface properties of a K12 *E. coli* bacterial strain that continuously expresses type 1 fimbriae. Figure 1a shows an electron micrograph of a typical bacterium having hundreds of fimbriae. Their contour lengths vary between a few hundred nanometers up to 2–3 micrometers (Figure 1b).

**Figure 1.**
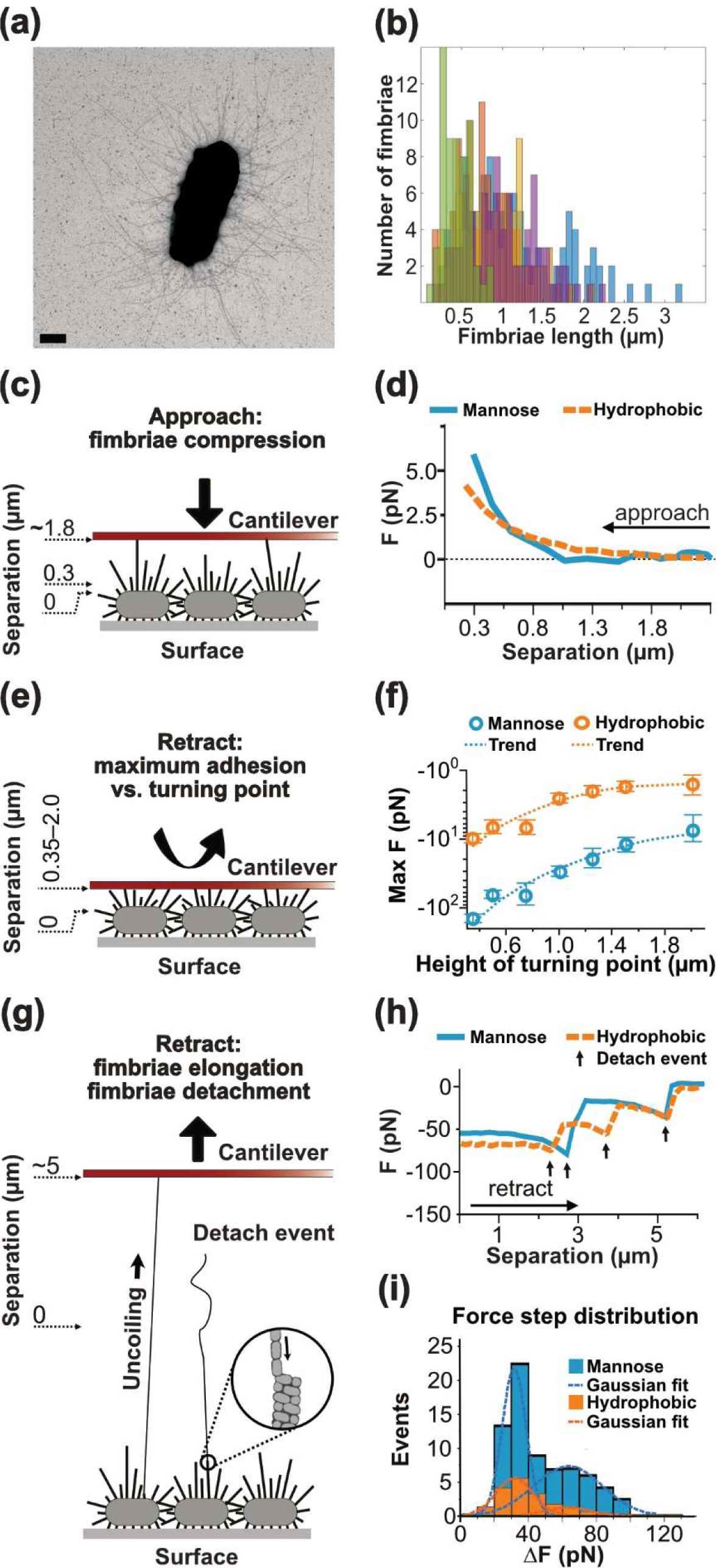
Characterization of the length distribution and load-bearing properties of fimbriae. (a) Transmission electron micrographs showing a fimbriated E. coli bacterium. The scale bar is 500 nm. (b) Histogram showing the contour lengths of the fimbriae of five different bacteria (colour-coded) obtained from analysis of TEM micrographs. (c) Illustration of how a flat AFM cantilever was brought into contact and pushed towards the fimbriae extending from bacteria immobilized side-by-side on a flat surface. (d) Typical repulsive force curves upon compression of the fimbriae extending from immobilized bacteria. The measured force was divided by the number of bacteria under the cantilever (30 for the hydrophobic cantilever and 10 for the mannosylated cantilever, respectively). (e) Illustration of the halting of the cantilever at different separations (0.35–2.0 µm) from the bacterial bodies before withdrawing. (f) The maximum adhesive force measured during retraction from the different turning separations. The measured force was divided by the number of bacteria under the cantilever (30 for the hydrophobic cantilever and 10 for the mannosylated cantilever, respectively). (g) The cartoon illustrates how the retraction of the cantilever forced attached fimbriae to uncoil (elongate) and eventually detach. (h) The graph shows details of retracting force curves emphasizing the last detachment events before complete cantilever detachment plotted on a common axis with approximate origin as indicated in the cartoon (g). (i) Histograms showing the distribution of all force steps recorded from the retract-curves using mannosylated and hydrophobic tips, respectively. The numbers were scaled by the number of bacteria under the cantilevers (30 for the hydrophobic cantilever and 10 for the mannosylated cantilever, respectively). Each step corresponds to the release of one or two fimbriae. Therefore, the data were fitted by two added Gaussians where the peak values of the first distribution, μ1, and the second distribution, μ2, were related through μ2=2μ1. This gave a peak value μ1=32.4±5.0 (R2=0.997) for mannose-coated cantilevers and μ1=32.7±7.5 (R2=0.996) for hydrophobic cantilevers, respectively.

AFM was used to probe the load-bearing properties of fimbriae upon binding specifically or non-specifically to tip-less (flat) gold-coated cantilevers modified with mannose-presenting or hydrophobic molecules, respectively (Figure 1c-i). The mannose coating was made by self-assembly of mannose-functional dendrimers with disulfide cores, previously shown to form monolayers on gold surfaces that have very high density of exposed mannose groups thus binding efficiently to type 1 fimbriated *E. coli*^56^. Fimbriae extending from approximately 10 surface-immobilized bacteria (when mannosylated cantilevers were used) or approximately 30 surface-immobilized bacteria (when hydrophobic cantilevers were used) were probed simultaneously (Supplementary figure 1). In each measurement cycle the cantilever was first lowered towards the bacteria until a threshold force of 0.6 nN was reached, therethrough defining the vertical position *z*=0 µm corresponding to the upper surface of the bacterial bodies, and then retracted to *z*=50 µm to ensure that all fimbriae were detached. The cantilever was then again approached towards the bacteria at constant rate (Figure 1c), but this time halted at a pre-set distance (*z*) above the bacterial bodies (Figure 1e), whereupon it was retracted until all fimbriae had detached (Figure 1g). Examples of full force-distance curves from these measurements are shown in Supplementary figure 2.

Upon approaching the cantilever, a repulsive force can be detected at separations smaller than approximately 1.5 µm from the bacterial bodies (Figure 1d). The repulsion increases monotonically with further decreasing separation. When subsequently retracting the cantilever, an adhesive force can be measured. The maximum force is reached after retracting the cantilever approximately 2 µm (Supplementary figure 2) and its magnitude varies inversely with the height *z* above the bodies of bacteria at which the cantilever had been halted (Figure 1f). Control experiments made with isogenic, *non-fimbriated* bacteria show that in absence of type 1 fimbriae on the bacterial surface, no interactions can be detected for *z*>0.2 µm. Thus, we considered the repulsive and adhesive forces detected for *fimbriated* bacteria at *z*>0.3 µm as solely due to interaction between fimbriae and the cantilever surface coating. This interpretation is supported by the fact that the retraction force-distance curves consist of several sequential plateaus and force steps (ΔF∼32 pN, Figure 1g-i) known to correspond to fimbriae elongation (uncoiling under constant force) and fimbriae detachment, respectively^20–22^.

The results obtained for the mannose-coated and hydrophobic cantilevers, respectively, are qualitatively similar, which indicates that the fimbriae’s quaternary structure does not collapse when they are compressed against a hydrophobic surface. Indeed, despite their large aspect ratio, type 1 fimbriae are known to be structures of extraordinary stability^57^. The persistence length of the type 1 fimbrial rod has been estimated to be about 50 µm, i.e., much longer than the fimbriae themselves^44^. Fimbriae will therefore provide resistance to their bending and compression, which explains why they can give rise to both repulsive and adhesive forces as a bacterium approaches a surface, as observed above. Under the simplified assumption that all (≍100) fimbriae between a bacterium and the cantilever contribute to the repulsive force measured for the full compression (to *z*=0.3 µm), the mean force per fimbria is ≍50 fN. Because of the fimbriae length distribution, more fimbriae can partake in repulsion and binding when the bacteria and the surface are forced into closer contact. Importantly, more long-lived fimbriae tethers to the surface form via specific fimbria–mannose binding than via non-specific binding to hydrophobic cantilevers at a given separation; this is observed from the order of magnitude larger maximum binding force per bacterium in contact with mannose-functionalized cantilevers (Figure 1f).

### Fimbriated bacteria change from transient to surface-mobile to immobile binding in response to increasing tether multivalency

The probability that a bacterium forms one or more fimbria tethers to a surface depends on the one hand on the specifics of the interaction and the number of fimbriae displayed by the bacterium. On the other hand, it depends on the density of binding sites on the surface. We patterned the walls of microfluidic channels with different surface densities of 10-nm hydrophobic or mannosylated gold nanoparticles (Supplementary Figure 3) to restrict and control the tether multivalency of the bacteria-surface interaction. The void area surrounding the nanoparticles was blocked for binding by grafting with poly(ethylene glycol), this way ensuring that each nanoparticle can only bind one fimbria^58^. We flowed bacteria through the channels at a low speed calculated to be ∼5 µm s^−1^ 1 µm above the surface plane (Supplementary figure 4). The surface-binding bacteria were recorded using bright-field microscopy; residence times ≥1 s at one position were registered as binding and used to compile heatmaps of bacterial attachment, and to calculate mean binding times (Figure 2). The corresponding distributions of binding times are shown in Supplementary Figure 5.

**Figure 2.**
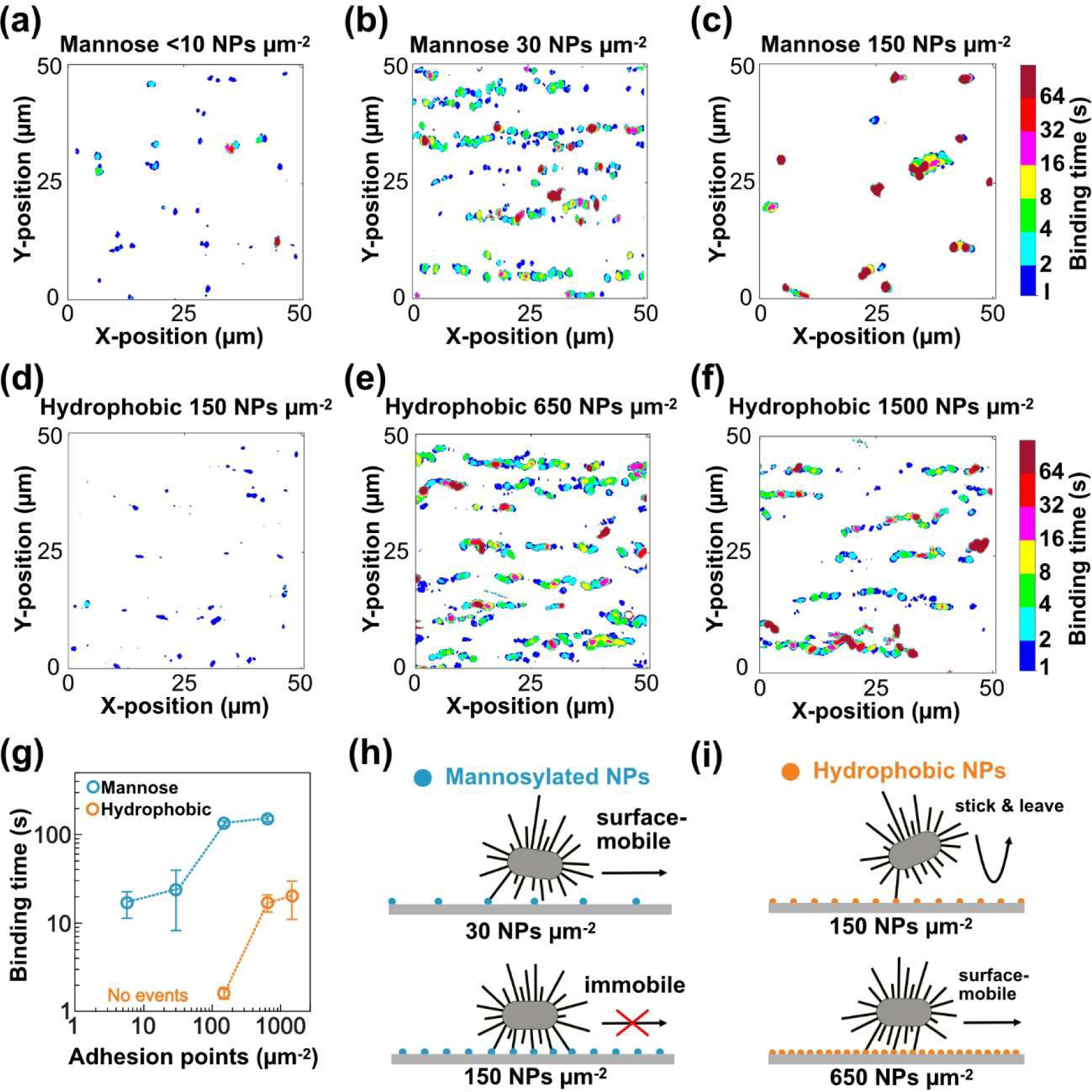
E. coli binding modes observed on surfaces for which the density and affinity of adhesion sites were varied. (a-f) Heat maps show the spatial distribution and binding time of bacterial footprints on microchannel surfaces displaying (a) <10, (b) 30, (c) 150 binding sites µm-2 of Au nanoparticles functionalized with mannosylated thiols or (d) 150, (e) 650 and (f) 1500 binding sites µm-2 of Au nanoparticles functionalized with hydrophobic thiols. The flow speed 1 µm above the surface was 5 µm s-1. (g) The graph shows the mean binding times (N≥3) plotted as a function of the surface concentration of binding sites (logarithmic scales). (h&i) Illustrations of the transition in binding behaviour observed in response to increased tether multivalency on surfaces with (h) mannose-modified (blue-coloured) and (i) hydrophobic (orange-coloured) adhesion sites.

Figure 2a-c displays binding of bacteria on surfaces coated with <10, 30, and 150 mannosylated adhesion points µm^−2^. The different surfaces gave rise to distinct binding patterns. On the surfaces with the lowest and intermediate densities of adhesion sites, bacteria bound transiently with similar mean binding times of ∼20 seconds (Figure 2a-b & g). However, while the lowest density of adhesion sites resulted in occasional “stick-and-detach” binding events (Figure 2a), the intermediate density enabled “stick-and-roll”^17^ and/or sliding surface motion of bound bacteria (Figure 2b). On surfaces with 150 adhesion sites µm^−2^, the binding times were much longer, and the displacements were, accordingly, rare (Figure 2c & g). The binding times remained unchanged upon further increase of the density of binding sites (Figure 2g).

The binding time of an *individual fimbria* depends merely on the specifics of the interaction and external forces acting on the bond^59^. Yet, the binding time of the *bacterium* will increase exponentially with the tether multivalency and therefore shift dramatically already for minor changes in the number of fimbria–surface bonds. A similar mean binding time (∼20 s) was detected in the experiments conducted with mannose-modified channels with <10 or 30 adhesion sites µm^−2^, indicating that the bacteria adhered through a similarly low number of fimbria tethers on these surfaces. As the few bond(s) break, the bacterium either leaves the surface (“stick-and-detach”) or makes a step along the surface (“stick-and-roll/slide”). The latter happens if the bacterium forms a downstream surface tether before or just after the initial upstream tether detaches. This interpretation corroborates the computer simulations by Whitfield *et al.*, which showed that surface motion is enabled whenever the rate of fimbriae binding is approximately balanced by the rate of fimbriae detachment^44^. Since the rate of bond formation is proportional to the density of adhesion sites under the bacterium, this requirement may be fulfilled by the increase from <10 to 30 adhesion sites µm^−2^ (Figure 2h). Upon a further five-fold increase to 150 adhesion sites µm^−2^, the surface-interacting bacteria will however form several additional tethers before the initially bound fimbria detaches. As a result, the equilibrium binding multivalency increases and the bacteria move much slower, i.e., become close to immobile on the surface (Figure 2h).

In contrast, on surfaces with the same density of adhesion sites (150 adhesion sites µm^−2^) but coated with *hydrophobic* molecules, only short (mean binding time 1.6 seconds) stick-and-detach binding events is seen (Figure 2d & g). A four-fold increase to 650 sites µm^−2^ led to a ten-fold increase in binding time and the transition from “stick-and-detach” to surface-mobile binding (Figure 2e & g) similar to that observed for bacteria binding to surfaces with 30 *mannosylated* adhesion sites µm^−2^. A further increase to 1500 hydrophobic adhesion sites µm^−2^ neither led to changed binding behavior nor increased binding times (Figure 2f & g). The non-specific bonds of fimbriae to hydrophobic surfaces are thus weaker than the specific bonds to mannosylated surfaces, in agreement with our AFM results. According to the discussion above, a large mismatch between the rate of bond formation and the rate of detachment of individual fimbriae should make stick-and-roll motion less likely. Yet, motion does occur on surfaces with a higher density (650 µm^−2^) of hydrophobic sites. This can be understood since already a minor rise in the density of adhesion sites triggers an increasingly multivalent binding and, thus, a steep, non-linear increase in binding time (Figure 2g & i). This result demonstrates that the concerted binding of several fimbriae can compensate for the low affinity that individual fimbriae have to hydrophobic surfaces and enable bacterial binding characteristics, including surface motion, similar to that observed for the specific FimH binding to mannose with much higher affinity. The fact that the binding time only increases slightly when the surface coverage of hydrophobic binding sites is raised to 1500 NPs µm^−2^ indicates that the equilibrium multivalency remains essentially unchanged, presumably because all available fimbriae on the bacteria already partake in binding.

### Surface-mobile E. coli slow down upon increased flow irrespective of the surface specificity

We also examined the response of bacteria that undergo surface motion when increasing the flow speed from ∼5 to ∼30 µm s^−1^ at 1 µm separation from the surface plane. On both mannosylated and hydrophobic surfaces, bacteria respond with stronger binding, the mean binding time increases from 20 seconds to 75 seconds and 63 seconds, respectively (Figure 3a-b and).

**Figure 3.**
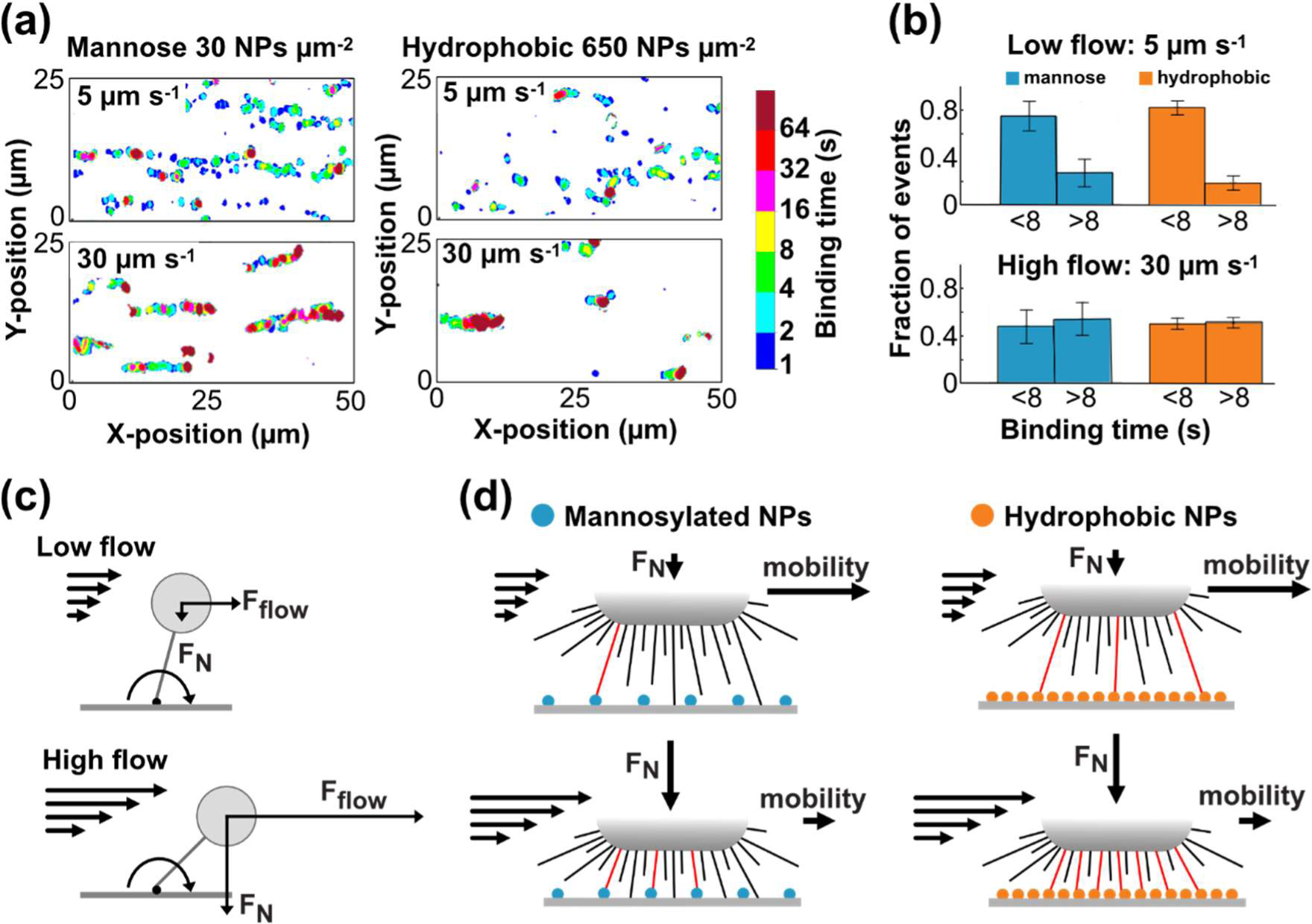
E. coli surface motion in response to an increased flow observed on surfaces where bacteria bound specifically and non-specifically. (a) Heatmaps show the spatial distribution and binding times of bacterial footprints on surfaces patterned with 30 mannose-modified adhesion sites µm-2 (left panels) and 650 hydrophobic adhesion sites µm-2 (right panels) before (upper panels) and after (lower panels) a flow increase from 5 µm s-1 to 30 µm s-1. (b) The histograms show the fraction of short (<8 s) and long (>8 s) binding times observed before and after the flow increase (N³3). (c-d) The cartoons illustrate the mechanism by which bacterial binding strength increases in response to increased flow force, Fflow. The flow force results in a torque on a tethered bacterium and a fraction (FN) of it pushes the bacterium closer to the surface, thus increasing the chance of multivalent binding irrespective of the surface specificity.

The force-enhanced binding of *E. coli* has been widely attributed to the catch-bond properties of the FimH–mannose interaction in combination with fimbriae extension due to fimbriae uncoiling^17,21,23,24^. Our observations cannot be explained by catch-bonds since we demonstrate the same flow-modulated binding behavior for bacteria binding to surfaces via non-specific fimbriae interactions. Moreover, activation of a catch-bond^18,19,60^ and fimbria uncoiling^20–22^ require high forces (typically >30 pN) acting along each surface-attached fimbria. In our experiments, the flow force acting on the bound bacterium as a whole after the flow increase is about 1 pN (Supplementary calculation). Our experiments performed at low constant flow featuring differently nanopatterned surfaces demonstrate that the bacterial binding time first increases drastically with the fimbriae tether multivalency but saturates for higher density of adhesion sites. The latter indicates that the number of fimbriae on a bacterium that can partake in binding becomes exhausted. Indeed, our AFM measurements (cf. Figure 1) show that in the absence of external forces other than the negligible gravity (corresponding to <10 fN^61^) the surface-approach of fimbriated bacteria may be limited due to fimbriae repulsion and the binding, accordingly, restricted to a few long fimbriae-surface tethers. A reasonable explanation for the observed flow-enhanced binding, which is also supported by previous theoretical work and simulations made by others^43,44^, is therefore that increased shear flow acting on a surface-tethered bacterium will produce torque and a normal force component strong enough to push it closer to the surface (Figure 3c). Thereby the number of fimbriae that can reach and form tethers to the surface, i.e. the multivalency of binding, will increase irrespective of the surface specificity (Figure 3d).

### E. coli are repelled from surfaces through the action of their longest fimbriae when the flow is low

To test the hypothesis of shear-flow driven force modulation of bacteria attachment via fimbriae tether valency, we investigated the flow-modulated surface motion of fimbriated and non-fimbriated bacteria with a modified version of digital holographic phase contrast microscopy^53^ (DHPCM). This method allows the three-dimensional position determination of all bacteria passing through microfluidic channels. As in previous experiments, the channel surfaces were modified with mannosylated or hydrophobic NPs to control the degree at which fimbriated bacteria interact with the channel walls. We also investigated, as a control, the behavior of non-fimbriated bacteria in the same channels. All bacteria flowing through the microfluidic channel were tracked continuously and their speeds along the flow direction, *v*_x_, were determined. Those traces originating from bacteria moving farther than 10 µm from the walls, the movements of which are unperturbed by surface interactions, were used to reconstruct the Poiseuille flow velocity profile across the channel (Figure 4a-b). Both the plane of the channel bottom, *z* = 0, (to within ±100 nm) and the local, instantaneous, flow speed, *v*_flow_(*z*), throughout the channel can be determined this way.

**Figure 4.**
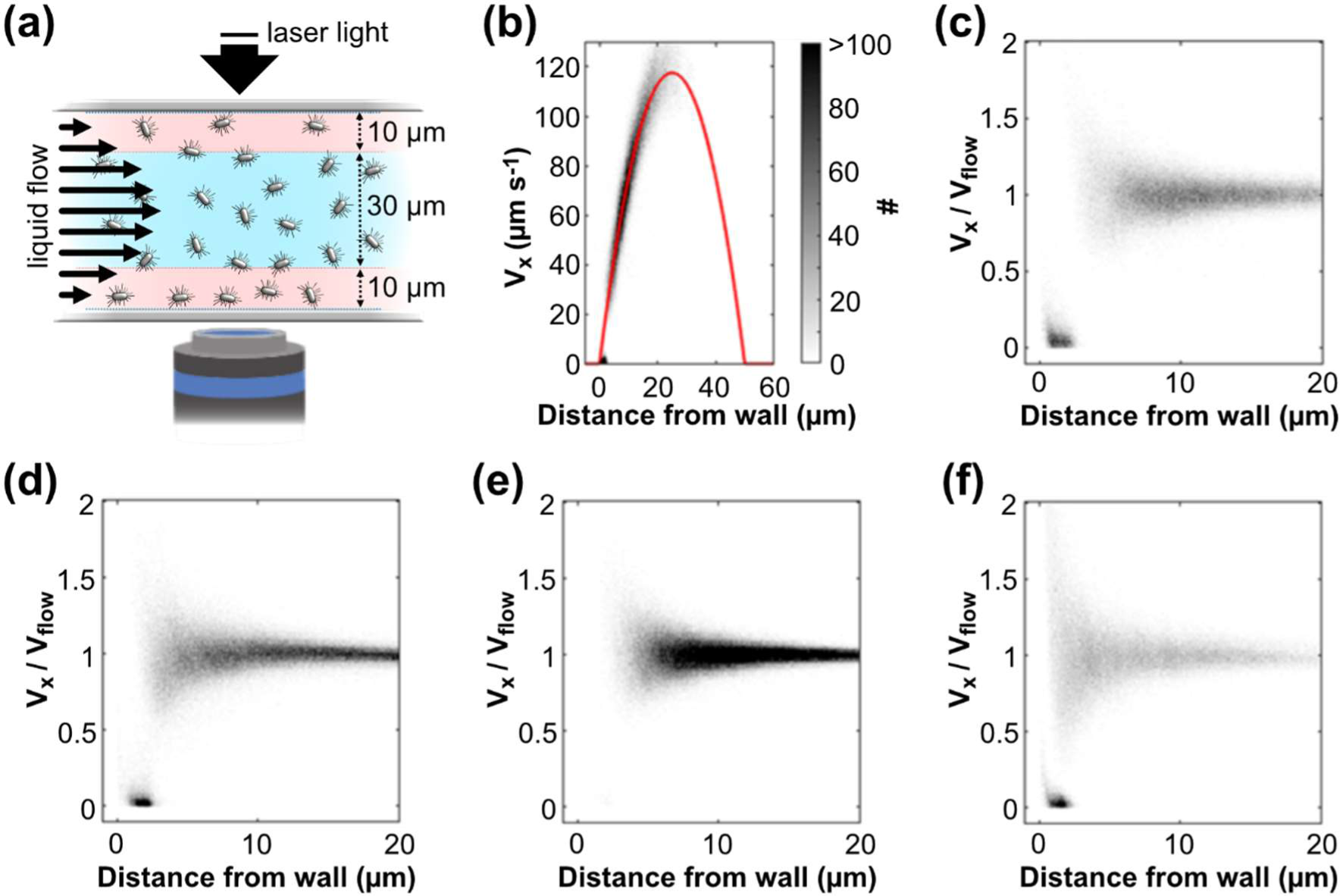
The 3D-distribution of fimbriated and non-fimbriated E. coli above surfaces with different adhesive properties. (a) The cartoon illustrates the setup used for digital holographic phase contrast microscopy. The movements of all bacteria passing through the channel were recorded, those originating from the central part of the channel were used to reconstruct the flow profile. (b) 2D histogram showing the distribution of the speed of bacteria, vx, flowing through the channel as a function of their separation from the channel’s bottom wall. The red line shows the Poiseuille flow velocity profile obtained from a fit to the bacterial speed data. (c) 2D histogram showing the distribution of the relative speeds, vx/vflow, of fimbriated E. coli flowing through a channel modified with 30 mannosylted NPs µm-2 as a function of their surface separation. (d) 2D histogram showing the distribution of the relative speeds, vx/vflow, of fimbriated E. coli flowing through a channel modified with 650 hydrophobic NPs µm-2 as a function of their surface separation. (e) 2D histogram showing the distribution of the relative speeds, vx/vflow, of fimbriated E. coli flowing through a channel modified with <10 hydrophobic NPs µm-2 as a function of their surface separation. (f) 2D histogram showing the distribution of the relative speeds, vx/vflow, of non-fimbriated E. coli flowing through a channel modified with 650 hydrophobic NPs µm-2 as a function of their surface separation.

We first studied the bacteria under a low flow determined to be approximately 10 µm s^−1^ at 1 µm separation from the surface. To identify the intermittent binding events that cause surface motion, it is instructive to study the ratio of the speed of a bacterium to the local flow speed at its position, *v*_x_⁄*v*_flow_, since surface binding will cause bacteria to slow down. Figures 4c-f show 2D histograms of *v*_x_⁄*v*_flow_ versus surface separation for all bacteria detected in different experiments featuring fimbriated bacteria on both types of surface coatings, and non-fimbriated bacteria on the hydrophobic coating, respectively. The speed of a suspended bacterium is the result of the local liquid flow speed at its position in the channel and its stochastic Brownian motion. Since the local flow speed is slow close to the surface, the distribution of relative speeds for the non-binding population therefore broadens with decreasing surface separation. For channels with coatings that support bacterial surface motion, all *fimbriated* bacteria found within 2-3 microns from the surface show strongly attenuated speed (Figure 4c-d). A similar zone, void of freely moving fimbriated bacteria, can also be seen in experiments featuring channels to which the bacteria did not bind (Figure 4e). In contrast, when *non-fimbriated* bacteria were flown through a channel coated with high surface concentration of hydrophobic NPs, we observe both surface-binding bacteria and numerous bacteria that move freely just a few hundreds of nanometers from the surface (Figure 4f). The non-fimbriated bacteria thus move with the flow, also with their bodies almost in contact with the surface since an *E. coli* body is typically 0.5–1 wide. The fimbriated bacteria are, however, separated from the surface by a distance that corresponds to that of their longest fimbriae. Notably, this is also just outside the region where fimbriae cause a repulsive force that can be measured with AFM (c.f. Figure 1d). This finding confirms our hypothesis that when the flow is low, the repulsive action executed by the longest fimbriae keeps the bacteria at such a large separation from the surface that few fimbriae reach the surface and partake in the initial binding. The resulting low binding valency explains why for weak individual fimbriae-surface bonds, like those with hydrophobic patches, bacteria will not be immobilized at low flow, even though the density of adhesion sites on the surface is high (c.f. Figure 2e&f).

### The longer fimbriae transform a flow increase into a compression

To scrutinize the influence of differently strong flow on the surface-binding bacteria, we analyzed the trajectories of individual fimbriated bacteria recorded in DHPCM experiments where the flow speed indicated at 1 µm surface separation was continuously increased from 5 to 100 µm s^−1^. Figure 5a displays the movements of a selected surface-interacting bacterium that could be traced throughout a large portion of the flow ramp. In the figure, the liquid flow speed, the speed of the bacterium, its position along the flow direction (*x*) and its position perpendicular to the surface plane (*z*) are plotted versus the same time axis, and the bacterium’s speed relative to the flow speed, *v*_x_⁄*v*_flow_, is indicated by the color of the trajectory.

**Figure 5.**
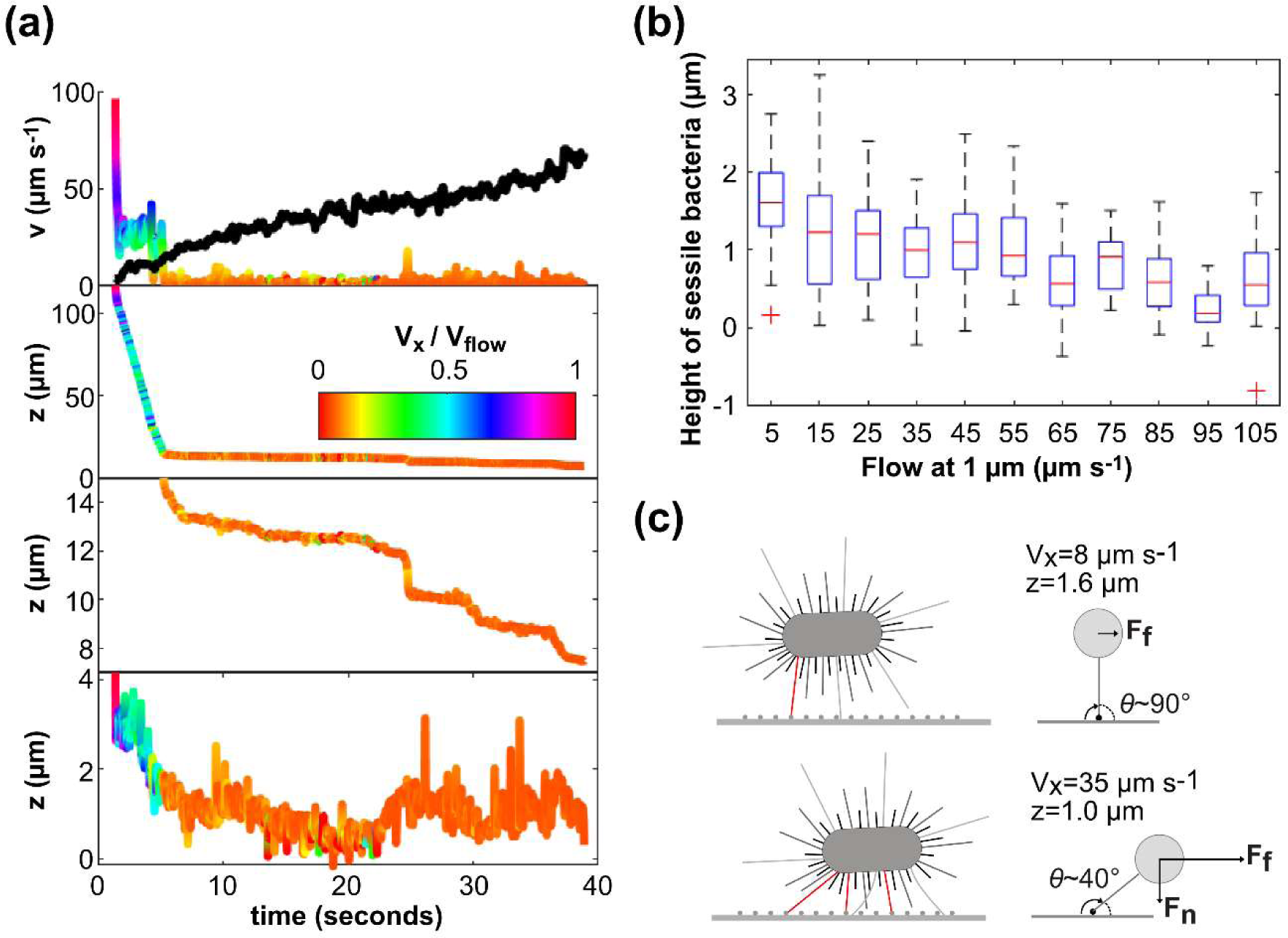
Influence of differently strong flow on the surface separation of bound fimbriated E. coli bacteria. (a) The combined plots show how the speed and movements of a selected fimbriated bacteria vary with time as the flow speed is continuously increased. The upper plot shows the flow speed measured at 1 µm surface separation (black line) and the speed of the bacterium, the center plots show the position of the bacterium on the axis parallel to the flow direction (x-axis), and the lower plot the position of the bacterium on the axis perpendicular to the channels bottom surface (z-axis). The bacterial trajectories are color coded according to the bacterium’s speed relative the local flow speed, *v*x⁄*v*flow. (b) The histogram shows the position above the surface plane where binding events (defined as a bacterium observation at *v*⁄*v*flow<0.25) were detected for all bacteria in a flow ramp experiment as a function of the flow speed in the channel. (c) A cartoon explaining how an increase in flow may result in a compressive force on a tethered bacterium and how the strength of this force may be estimated in this situation.

After binding, which happens early during the flow ramping procedure at t≍5 s, the bacterium displays the characteristic surface motion observed in the previous experiments. This encompasses shorter periods of faster step-wise movements separated by longer periods of much slower, creeping motion, although the bacterium is hardly ever completely still. Careful inspection of the trajectory reveals that each period of faster movement is preceded by the bacterium shifting position slightly away from the surface plane. During the movements forward, it instead approaches the surface, i.e., the overall movement is diagonally downwards. The intervening creeping motion, in contrast, correlates to periods characterized by less vertical motion of the bacterium. The accumulated effect of the diagonal movements following on each other is that the bacterium relocates from a position almost 2 µm above the surface plane in the beginning of the flow ramp, to below 1 µm surface separation halfway through the flow ramp at t≍20 s. This point of closest approach also correlates with the slowest motion and the strongest binding, i.e., the slowest relative speed of the bacterium. To verify that the observed effect of increased flow speed on the bacterium’s vertical position indeed is a general phenomenon, we accumulated statistics for this for all bacteria present during the flow ramp experiment and plotted the data as a function of flow speed (Figure 5b). Here, binding events were defined as bacterial positions for which *v*_x_⁄*v*_flow_ < 0.25; the DHPCM data acquired at low flow speed (c.f. Figure 4c-d) show that this cutoff value can separate bond from non-bond bacteria reasonably well. The boxplot of Figure 5b reproduces the main characteristics of the single-bacterium trajectory; bacteria displace downward as the flow speed increases reaching a maximum shift of approximately 1 µm. The bacterium highlighted in Figure 5a however responds stronger to already a smaller flow increase than most of the bacteria detected in the flow ramp experiment do. In the general case, the smallest surface separation was seen towards the end of the flow ramp when the flow rate was higher than 50 µm s^−1^.

We interpret the repeated diagonal displacement towards the surface observed in the single-bacterium trajectory as the rotational motion of a bacterium pivoting around one of its surface-tethered fimbria. Due to the tethering, a hydrodynamic drag force (F_F_), the magnitude of which is proportional to the difference between the bacterium’s velocity and the speed of the surrounding liquid flow, acts on the bacterium in the direction of the flow. We estimate the proportionality constant to 1.9 × 10^−8^ kg × s^−1^ (Supplementary calculation). In addition, the tethered bacterium experiences a torque and, consequently, a force normal to the surface plane (F_N_) that pulls it downwards towards the surface (Figure 5c). The magnitude of F_N_ depends both on F_F_ and the angle between the attached fimbria and the surface. As F_F_ increases so does F_N_ and eventually it overcomes the repulsive force exerted by the longer fimbriae, bringing the bacterium closer to the surface. The approach will, however, again be halted due to the balancing stronger repulsive force emerging from a larger number of shorter fimbriae that are brought into surface contact and become compressed (Figure 5c).

Based on this model, we may use the first part of the DHPCM flow ramp data in combination with the AFM results to estimate the sequence of events underlying the bond strengthening observed in the bright-field microscopy experiment above (c.f. Figure 3). As the flow rises from 5 to 30 µm s^−1^, the bacteria typically displace from 1.6 to 1.0 µm above the surface (Figure 5b) and F_F_ simultaneously increases from 0.15 pN to 0.6 pN (Supplementary calculation). If the initial attachment takes place through a long fimbria binding with a nearly ninety-degree angle relative the surface and the bacterium pivots around this attachment point, the corresponding F_N_ that push the bacterium closer to the surface is on the order of 0.3 pN (Figure 5c and Supplementary calculation). The magnitude of the repulsive barrier preventing bacteria from strong binding at low flow thus correlates with force needed to compress a handful of fimbriae as determined by scaling with the estimated repulsive force per fimbria measured by AFM. This is a reasonable number also considering the typical distribution of long versus short fimbriae revealed by electron microscopy (Figure 1b). The large number of shorter fimbriae that come within reach of the surface not only provides a second stronger repulsive barrier to further approach, but it also increases the probability that additional fimbriae bind to the surface. AMF retraction data suggests that approximately twice as many bonds form when the separation between bacteria and the surface decreases by 0.6 µm (c.f. Figure 1f). Given that the bacteria’s binding time is expected to depend exponentially on the number of tethers, this difference may readily explain the decrease in bacterial surface-mobility by a factor 3-4 observed when the flow was increased from 5 to 30 µm s^−1^ (c.f. Figure 3).

The impact of the flow force on F_N_ and, in turn, the impact of F_N_ on the bacteria’s position beyond the initial binding and approach event is more ambiguous and therefore less straightforward to analyse. Bond(s) established between the shorter fimbriae and the surface will maintain the bacterium in proximity of the surface, also if flow forces vanish. The binding configuration is, however, not static as evidenced by the fact that surface-attached bacteria move along the surface through repeated small steps, which are due to the process of fimbriae continuously binding and unbinding: When a load-carrying surface-attached fimbria releases, the bacterium moves slightly until other fimbriae become strained.

Fimbriae assumingly bind and unbind independently from each other; thus, the frequency of unbinding events leading to redistribution of adhesion forces is expected to increase with the multivalency of the binding. In this regime the small details of the geometry of binding, and so the magnitude of F_N_, may fluctuate rapidly and stochastically. If, because of the changing binding configuration, the most load-carrying fimbria is comparably long and its binding angle either is very low (fimbria close-to aligned with surface) or very high (θ>90°), F_N_ will drop. These types of events thus explain the occasionally increasing surface separation observed in the single-bacterium trajectory, typically preceding the downward diagonal movements. Notably, the equilibrium bacteria-surface separation observed for an applied flow, and thereto related binding behavior, will be sensitive to the number-length distribution of fimbriae on a bacterium. This may vary considerably between different bacteria (c.f. Figure 1b) with some bacteria having only short fimbriae while others having additional extra-long ones, explaining the variability of the data presented in Figure 5.

## Discussion

We show that the intrinsic capacity of multivalent binding enables fimbriated bacteria to switch between transient, rolling, sliding, and immobile binding modes. Since this type of regulation through tether multivalency determines a sharp response already to minor differences in the number of binding sites on a surface as well as the number and flexibility of the tethers that mediate binding, it is often referred to as “super-selective”^42^. We show that the fimbria-mediated surface repulsion and the flow pressure acting on a bacterium establish an equilibrium bacterium–surface separation that dictates the maximum number of fimbriae that can reach the surface. For a constant density of surface ligands, the threshold flow rate at which the switch between transient and strong biding modes occurs is tuned by the mechanical stiffness and affinity of the individual fimbriae and their number/length distribution on the cell surface. We show that this mechanism *per se* is not dependent on the type of bond formed by the fimbriae to the surface. Accordingly, this kind of flow/force-modulated binding can exist for any bacteria/cell/particle that binds to surfaces via multiple elastic extensions.

The type 1 fimbriated *E. coli* adapt their avidity to the environmental settings, in contrast to bacteria that lack adhesion organelles and attach to surfaces via contact points^62^ or patches^63^ distributed directly on the bacterial body. We show that the combination of weak individual fimbriae–surface bonds, stiff elastic extensions, and low flow allows these bacteria to bind to and remain surface-associated (rolling/sliding), while the extensively multivalent, strong binding state is prohibited. Therefore, it is interesting that *E. coli* carrying mutations of FimH that increase the strength of individual fimbriae–mannose bonds^11,64^ or a mutation of FimA that decreases the mechanical stability of the fimbrial shafts^37^ have proven deficient in infecting the urinary tract of mice. Other studies have shown that fimbriae are not needed for bacteria to adhere persistently to abiotic surfaces but are necessary for biofilms to develop^38–40^. Furthermore, we and others^19,65^, observe that the transition from surface-mobile to stable binding occurs for flow forces (in the range of 1 pN) that are physiologically ubiquitous. The emerging picture is thus that the flow-modulated binding mechanism discussed in this work is not primarily a protection against extreme flows. Dealing with the latter may be the specific function of catch-bonds and fimbriae uncoiling that occur at forces >30 pN per attached fimbria for their activation. In contrast, fimbriae’s main advantage appears to be associating bacteria to surfaces without immobilizing them in naturally occurring weak flow. This can, e.g., aid the expansion of growing microcolonies^65^ or direct bacteria from unfavorable to better binding positions^66^ in a heterogeneous microenvironment. The ecological impact of surface-sensing and exploration mechanisms for successful surface colonization has been established for *Pseudomonas aeruginosa*^67–69^ and *Vibrio Cholerae*^70^.

This study establishes that the elastic type 1 fimbriae, and potentially other CU pili, provide *E. coli* with a *generic* mechanism to sense and adapt their binding behavior to different surface conditions. In contrast to the widely-spread interpretation that fimbriae’s function is determined by their lectins, we show that force-modulated binding is not limited to certain types of specific binding and may benefit from individual fimbria–surface bonds being weak. This framework is essential to understand how fimbriae/pili contribute to bacterial tropism and the process of biofilm formation on both living tissue and artificial surfaces such as catheters. We expect it to apply also to other surface-colonizing types of cells with elastic surface appendices.

## Material and methods

### Bacterial cultures

Isogenic mutants of *E. coli* K12 strain PC31 that lack type 1 fimbraie (PC31 Δ*fim* Km^r^)^71^ or express type 1 fimbraie continously (PC31 Δ*fim* Km^r^ pPKL4)^71^ were used for all experiments. The plasmid pPKL4 encodes all *fim* genes and resistance to ampicillin^72^. The expected phenotypes were confirmed by electron microscopy (TEM) and a yeast-agglutination assay. TEM analysis and the observation of few swimming bacteria with light microscopy showed that only a small fraction of the bacteria had flagella. The bacteria were stored in deep-freeze cultures and cultivated on Lauria-Bertani (LB) - Agar plates supplemented with appropriate antibiotics. New plates were streaked once a week, cultivated for 24 h at 37°C and then stored at 4–8 °C. Bacteria solution for experiments was prepared by inoculating a single colony in 15 ml of LB media, supplemented with appropriate antibiotics, in a 50 ml Falcon tube which was then incubated overnight (o.n.) at 37 °C under stirring conditions. Type-1 fimbriated *E. coli* are prone to form irreversible aggregates when growing in liquid culture^73^. To get a solution containing mainly single bacteria, aggregates were separated by centrifugation of the o.n. culture at 1,000 × *g* for 4 minutes. The supernatant was diluted with fresh LB media or PBS buffer pH 7.0 to the concentration that was suitable for the experiment.

### Electron microscopy (EM)

EM grids coated with a 3–4 nm thick carbon film (CF300-CU-UL, Electron Microscopy Sciences, USA) were treated in an UV/ozone chamber (ProCleaner, Bioforce Nanoscience, USA) for 1 minute and then put up-side-down on a droplet of bacterial solution with optical density (OD, measured at λ=600 nm) >0.5 for 1 minute to allow bacteria to adsorb. The adsorbed material was fixed with glutaraldehyde (2.5% w/w solution in PBS buffer pH 7.4) for 10 minutes, rinsed 3×10 seconds with MQ water and then negatively stained with uranylacetate (1% w/w solution in MilliQ water) for 1 minute. Electron micrographs were captured using a FEI Tecnai G2 microscope operated at 160 kV acceleration voltage. The contour lengths of fimbriae adsorbed on the surface, surrounding different bacteria were measured manually using the open-source software ImageJ^74^.

### Atomic Force Microscopy (AFM)

Sample surfaces with immobilized bacteria were prepared on circular (ø = 24 mm) glass coverslips. The slides were cleaned first by immersion in Hellmanex III solution (2% v/v, Hellma Gmbh, Germany) o.n. and then in 2 M H_2_SO_4_ (aq.) for 30 minutes. After rinsing first with MQ water and then with methanol (analytical grade) the glass slides were carefully dried under a flow of N_2_ gas and then immersed in a 5% v/v methanol solution of (3-aminopropyl)dimethylethoxysilane (APDMES 95%, ABCR GmbH, Germany) for 30 minutes. After rinsing first with methanol and then with MQ water, the silanized slides were reacted with glutaraldehyde (GA, 2.5 % v/v solution in PBS buffer pH7.4) for 30 minutes. Bacteria were cross-linked to the GA-activated surfaces by immersing them in bacteria solution diluted to OD ∼0.2 in PBS for 5 or 30 minutes to make surfaces with low and high bacteria coverages, respectively (Supplementary figure 1). Non-reacted GA was then blocked by reaction with Bovine Serum Albumin (BSA, 1 mg/ml in PBS buffer) for 30 minutes. Sample surfaces were kept at 4–8 °C until use within 24 h.

Tipless gold-coated cantilevers with nominal spring constant 0.06 N/m were cleaned in a UV/Ozone chamber (ProCleaner, Bioforce Nanoscience, USA) and surface modified by immersion either in 1 mM ethanolic solution of 1-Octanethiol or 100 µM ethanolic solution of Mannose-functional first-generation dendrimer with disulfide core (G1-SS-Man4-sp). The synthesis and application of G1-SS-Man4-sp for the self-assembly of mannose-presenting gold surfaces are detailed in Öberg *et al*^56^. AFM was done with a JPK Nanowizard III (Bruker Nano GmbH, Germany) with the CellHesion module and JPK’s custom thermo-regulated flow-cell (JPK Instruments, Germany) mounted on an inverted optical microscope (Axio Observer Z1, Zeiss, Germany) to allow precise positioning of the cantilever over groups of immobilized bacteria. The samples were kept in PBS at 37 °C throughout the measurements. The AFM was operated in force-spectroscopy mode, the approach/retract speed was held constant at 2 µm s^−1^ with minimum delay (<0.5 s) between approach and retract phases. In each measurement cycle the cantilever was first lowered towards the bacteria until a threshold force of 0.6 nN was reached, therethrough defining the vertical position *z*=0 µm corresponding to the upper surface of the bacterial bodies, and then retracted to *z*=50 µm to ensure that all fimbriae were detached. Approach/retract force curves were recorded while bringing the cantilever to different turning points *z* =0.1–2 µm above the bacteria. In between fimbriae capture experiments, the glass substrate was probed to ensure cantilever cleanliness. Experiments with *non-fimbriated* bacteria occasionally showed weak adhesive force for *z* <0.2 µm. Thus, we considered adhesive force detected for *fimbriated* bacteria at *z* ≥0.35 µm as due to fimbarie attachment. Data were processed using JPKSPM software (JPK Instr., Germany).

### Microfluidic channels

Microfluidic channels for the bright-field microscopy (BF) and digital holographic phase contrast microscopy (DHPCM) were assembled from rectangular Borosilicate glass capillaries (VitroCom Inc., USA). Capillaries with inner dimensions 0.8×0.8 mm and wall thicknes 0.16 mm were used for BF and capillaries with inner dimension 0.05×1.0 mm and wall thickness 0.05 mm were used for DHPCM. To enable their mounting on the microscope stage, the capillaries were glued to supporting glass sildes. Tubings was connected to a capillary by inserting very narrow pipette tips (200 µl gel-loading tips, VWR) into its ends and fixating these/sealing the void space with a UV-curable glue (NOA68, Norland Products Inc.,USA). The inner walls of the capillaries were provided chemical nanopatterns by the means of Debye screening-controlled nanoparticle (NP) self-assembly, previously outlined and used for analysis of cell and bacteria binding by Lundgren *et al.*^58,75^. Firstly, the insides of the glass capillaries were cleaned and modified with APDMES as detailed for AFM sample surfaces above. Secondly, gold NPs with an approximate diameter of 10 nm were synthesized by reduction of HAuCl_4_ with a mixture of sodium citrate and tannic acid as reductive agents^76^. Briefly, one solution of 80 ml of 0.32 mM HAuCl (Aldrich, 99.999%) in ultra-pure water and another solution of 20 ml containing 6.8 mM sodium citrate (Sigma, >99.0%) and 0.24 mM tannic acid (Sigma-Aldrich) in ultra-pure water were both heated to 60°C. The two solutions were then mixed and allowed to react for 25 minutes while maintaining the temperature at 60°C whereupon the mixture was quickly heated to 95°C. The final concentration of NPs was estimated to approximately 10 nM based on the assumption that all gold ions had been reduced to gold. The NPs were centrifuged twice to remove excess sodium-, citrate- and tannate ions from the solution and to increase the NP concentration. Arrays with decreasing density of gold NPs were deposited onto the amino-silanized surfaces by incubating these 15 minutes under static conditions with NPs solved in citrate buffer pH 4.0 (CB) with decreasing ionic strength and/or concentration. The following conditions were used: (i) 30 nM NPs in 10 mM CB, (ii) 30 nM NPs in 0.08 mM CB, (iii) 3 nM NPs in 0.08 mM CB, (iv) 0.3 nM NPs in 0.08 mM CB, (v) 0.03 nM NPs in 0.08 mM CB. The resulting NP coverages were estimated from electron micrographs by performing the same NP modifications on TEM grids coated with SiO (SF300-CU, Electron Microscopy Sciences, USA) as shown in Supplementary figure 3. Thirdly, after rinsing with MQ and PBS buffer pH 7.4, amino groups of APDMES not blocked by NPs were reacted with1 mM methoxypolyethylene glycol 5,000 propionic acid N-succinimidyl ester in PBS pH 7.4 o.n. at 4–8 °C^77^. Finally, just before use, the capillaries were rinsed with MQ and Ethanol whereupon the surfaces of the gold NPs were modified by recation with 1 mM etanolic solution of 1-Octanethiol or 0.1 mM ethanolic solution of G1-Mannose^56^ for 30 minutes.

### Bright-field microscopy (BF) experiments

BF microscopy was performed using an up-right microscope equipped with a 60× water-immersion objective with NA=1.0 and 2 mm working distance that allowed focusing on the bottom of the microfluidic channel. The channels were rinsed with MQ water and LB-media whereupon bacteria solution diluted to OD=0.2 in LB media was injected at flow rate 16 µl/min or 100 µl/min using a syringe pump (in-house built). The corresponding flow speeds at 1 µm surface separation was calculated according to Lima *et al.*^78^ (Supplementary figure 4). Digital micrographs (8-bit greyscale) were acquired using 1 second exposure time at 1 fps for periods of 600 seconds. The image sequences were treated and analyzed using methods and scripts for ImageJ and Matlab. This involved, in following order, (i) correction for uneven illumination by “rolling ball” background subtraction, (ii) correction for drift by xy-alignment of sequential images, (iii) thresholding with manually selected threshold and, (iv) removal of noise by median filtering. The intensity threshold was set to keep only those pixels that were occupied by a bacterial footprint during the full exposure. Heatmaps showing the position and duration of binding were constructed by setting the value of the occupied pixels in each image to 1 second and then summing all images in a sequence. The distribution of binding times and mean binding time were calculated based on the pixel values of the heatmaps.

### Digital holographic phase contrast microscopy (DHPCM) experiments

DHPCM experiments were conducted on inverted microscope (Nikon eclipse TS 100) equipped with a Plan Apo 60× water-immersion objective with a Numerical Aperture of 1.3 and a correction ring to correct for the spherical aberration associated with a coverslip. The sample was illuminated by the beam of a laser pointer (405nm, 5mw, teknistore.com) replacing the lamp and condenser lens for Köhler illumination. The focal plane of the microscope was set a few tens of micrometers below the channel bottom. The channels were rinsed with MQ water and LB-media whereupon bacteria solution diluted to OD=0.2 in LB media was injected using a syringe pump. A low flow rate resulting in a flow speed of 5– 10 µm/s at 1 µm surface separation was applied for 600 seconds whereupon the flow was linearly increased. Digital micrographs of the interference between light scattered by the bacteria and the unscattered illuminating beam were acquired using 1 ms exposure at 25 fps of 12 bits greyscale images, using a CMOS camera (Basler MK3 acA2000-340km), in combination with a Solios 2M evCLF framegrabber, operated using an in-house GUI based on the Image Acquisition Toolbox of Matlab. The image sequences were saved as uncompressed .mj2 movie files. The images were filtered with a Gaussian filter with a standard deviation of 64 pixels to be used as a background by which the original images are divided. This procedure removes low spatial frequency variations in the illuminating intensity but importantly retains the fringes associated with non-motile bacteria, in contrast to the more commonly used time averaged background that also removes fringes associated to dust and scratches on optics on nearby (conjugate) planes. These background-corrected images are backpropagated using the Rayleigh-Sommerfeld propagator to generate a 3D reconstruction of the instantaneous scattered light field. In this lightfield we isolate voxels that are associated to bacteria using a modified version^79^ of the method of Cheong *et al.*^53^. This method yields an improved contrast with respect to that of the scattered light by considering the phase of the scattered light and shifting it by Ø, chosen to be π, such that the background signal is next to zero. Essentially, we numerically generate a stack of images similar to what would be obtained in the normally dark arm of a Mach-Zehnder interferometer. The data is thresholded both on the intensity and the phase of the scattered light^79^ and labeled to form 3d images of the individual bacteria of which the positions are estimated as the intensity weighted average of the associated voxel positions. Repeating this analysis on a time-series of frames allows us to track the bacteria as they move through space. The poiseuille flow in a rectangular capillary is fitted using a Levenberg-Marquardt algorithm to the apparent displacements and simultaneously guides the search for candidate movements^80,81^, which is particularly useful to stitch time series of individual bacterium positions together at elevated flowrates. From the covariance of the fit of the flow profile after disgarding outliers as well as any bacteria closer than 10 µm to the wall we conclude that we find the position of the no-slip boundary layer to within 50 nm accuracy. The bacterium-wall interactions cause the bacteria to occasionally slow down or accelerate with respect to the local Poiseuille flow flow. The bacterium trajectories are determined by first expanding developing trajectories with candidates around the position they would be at if they continued to move the way they were moving, then by looking around the flow displaced position and finally around the position in the previous frame. Any bacteria not assigned to ongoging trajectories start a new one. We consider a bacterium ‘surface active’ if it is moving less than 25% of the flowspeed at its height.

## Supplementary information

Supplementary figure 1: Samples with immobilized *E. coli* used for AFM experiments

Supplementary figure 2: Full approach-and-retract force spectra obtained with AFM

Supplementary figure 3: Surface concentration of gold nanoparticles obtained for different adsorption conditions visualized by electron microscopy

Supplementary figure 4: Calculated flow profiles in the microfluidic channels used for bacterial binding experiments recorded with brightfield microscopy

Supplementary figure 5: Distribution of binding times measured for *E. coli* binding to different nanopatterns determined from footage acquired with brightfield microscopy Supplementary calculation: Estimation of forces acting on surface tethered bacteria subject to flow

## Acknowledgements

Professor Malte Hermansson is acknowledged for providing the bacterial strains used in the study. Dr. Andrea Lassenberger is acknowledged for assistance with TEM acquisition of gold nanoparticle patterns. Andrea Scheberl is acknowledged for assistance with sample preparations for bacteria TEM. The research leading to these results received funding from the European Union (MSCA-IF 752175), from the European Research Council under the European Union’s Seventh Framework Programme (FP/2007−2013)/ERC Grant Agreement 310034, and from the Swedish Scientific Council (Grant 2019-05215).

## Author contribution statement

**AL** conceptualized the study, made the experimental work involving surface chemical modifications, microbiological procedures, brightfield and DHPCM data acquisition, analysis of brightfield data, acquisition and analysis of bacteria TEM data, took part in planning and analysis of AFM experimnets and wrote the manuscript draft. **PvO** conceptualized the study, developed and implemented the DHPCM method, took part in DHPCM data acquisition, analyzed all DHPCM data and wrote the manuscript draft. **JI** planned, implemented, and analyzed the AFM experiments. **JLT-H** took part in planning of AFM experiments and the analysis of AFM data. **MM** synthesized the G1-SS-Man4-sp molecules used to create mannosylated nanopatterns. **ER** conceptualized the study and took part in the data analysis and writing of the manuscript draft. All authors took part in finalizing the manuscript.

## References

1. Waksman, G. & Hultgren, S. J. Structural biology of the chaperone–usher pathway of pilus biogenesis. Nat. Rev. Microbiol. 7, 765–774 (2009).

2. Wurpel, D. J., Beatson, S. A., Totsika, M., Petty, N. K. & Schembri, M. A. Chaperone-Usher Fimbriae of Escherichia coli. PLOS ONE 8, e52835 (2013).

3. Mulvey, M. A. Induction and Evasion of Host Defenses by Type 1-Piliated Uropathogenic Escherichia coli. Science 282, 1494–1497 (1998).

4. Connell, I. et al. Type 1 fimbrial expression enhances Escherichia coli virulence for the urinary tract. Proc. Natl. Acad. Sci. 93, 9827–9832 (1996).

5. Roberts, J. A. et al. The Gal(alpha 1-4)Gal-specific tip adhesin of Escherichia coli P-fimbriae is needed for pyelonephritis to occur in the normal urinary tract. Proc. Natl. Acad. Sci. 91, 11889–11893 (1994).

6. Korhonen, T. K. et al. Binding of Escherichia coli S fimbriae to human kidney epithelium. Infect. Immun. 54, 322–327 (1986).

7. Jordan, D. M. et al. Long Polar Fimbriae Contribute to Colonization by Escherichia coli O157:H7 In Vivo. Infect. Immun. 72, 6168–6171 (2004).

8. Sakellaris, H., Munson, G. P. & Scott, J. R. A conserved residue in the tip proteins of CS1 and CFA/I pili of enterotoxigenic Escherichia coli that is essential for adherence. Proc. Natl. Acad. Sci. 96, 12828–12832 (1999).

9. Barbercheck, C. R. E., Bullitt, E. & Andersson, M. Bacterial Adhesion Pili. in Membrane Protein Complexes: Structure and Function 1–18 (Springer Nature Singapore, 2018).

10. Korea, C.-G., Ghigo, J.-M. & Beloin, C. The sweet connection: Solving the riddle of multiple sugar-binding fimbrial adhesins in Escherichia coli: Multiple E. coli fimbriae form a versatile arsenal of sugar-binding lectins potentially involved in surface-colonisation and tissue tropism. BioEssays 33, 300–311 (2011).

11. Kalas, V. et al. Evolutionary fine-tuning of conformational ensembles in FimH during host-pathogen interactions. Sci. Adv. 3, e1601944 (2017).

12. Klemm, P. & Christiansen, G. Three fim genes required for the regulation of length and mediation of adhesion of Escherichia coli type 1 fimbriae. Mol. Genet. Genomics 208, 439–445 (1987).

13. Sharon, N. Bacterial lectins, cell-cell recognition and infectious disease. FEBS Lett. 217, 145–157 (1987).

14. Brinton, C. C. The structure, function, synthesis and genetic control of bacterial pili and a molecular model for DNA and RNA transport in Gram negative bacteria. Trans. N. Y. Acad. Sci. 27, 1003–1054 (1965).

15. Klemm, P. The fimA gene encoding the type-1 fimbrial subunit of Escherichia coli. Nucleotide sequence and primary structure of the protein. Eur. J. Biochem. 143, 395–399 (1984).

16. Thomas, W. E., Trintchina, E., Forero, M., Vogel, V. & Sokurenko, E. V. Bacterial Adhesion to Target Cells Enhanced by Shear Force. Cell 109, 913–923 (2002).

17. Thomas, W. E., Nilsson, L. M., Forero, M., Sokurenko, E. V. & Vogel, V. Shear-dependent ‘stick-and-roll’ adhesion of type 1 fimbriated Escherichia coli: Stick-and-roll bacterial adhesion. Mol. Microbiol. 53, 1545–1557 (2004).

18. Yakovenko, O. et al. FimH Forms Catch Bonds That Are Enhanced by Mechanical Force Due to Allosteric Regulation. J. Biol. Chem. 283, 11596–11605 (2008).

19. Aprikian, P. et al. The Bacterial Fimbrial Tip Acts as a Mechanical Force Sensor. PLoS Biol. 9, e1000617 (2011).

20. Miller, E., Garcia, T., Hultgren, S. & Oberhauser, A. F. The Mechanical Properties of E. coli Type 1 Pili Measured by Atomic Force Microscopy Techniques. Biophys. J. 91, 3848–3856 (2006).

21. Forero, M., Yakovenko, O., Sokurenko, E. V., Thomas, W. E. & Vogel, V. Uncoiling Mechanics of Escherichia coli Type I Fimbriae Are Optimized for Catch Bonds. PLoS Biol. 4, e298 (2006).

22. Andersson, M., Uhlin, B. E. & Fällman, E. The Biomechanical Properties of E. coli Pili for Urinary Tract Attachment Reflect the Host Environment. Biophys. J. 93, 3008–3014 (2007).

23. Zakrisson, J., Wiklund, K., Axner, O. & Andersson, M. The Shaft of the Type 1 Fimbriae Regulates an External Force to Match the FimH Catch Bond. Biophys. J. 104, 2137–2148 (2013).

24. Whitfield, M. J., Luo, J. P. & Thomas, W. E. Yielding Elastic Tethers Stabilize Robust Cell Adhesion. PLoS Comput. Biol. 10, e1003971 (2014).

25. Flores-Mireles, A. L., Walker, J. N., Caparon, M. & Hultgren, S. J. Urinary tract infections: epidemiology, mechanisms of infection and treatment options. Nat. Rev. Microbiol. 13, 269–284 (2015).

26. Eris, D. et al. The Conformational Variability of FimH: Which Conformation Represents the Therapeutic Target? ChemBioChem 17, 1012–1020 (2016).

27. Cusumano, C. K. et al. Treatment and Prevention of Urinary Tract Infection with Orally Active FimH Inhibitors. Sci. Transl. Med. 3, 109ra115–109ra115 (2011).

28. Han, Z. et al. Lead Optimization Studies on FimH Antagonists: Discovery of Potent and Orally Bioavailable Ortho-Substituted Biphenyl Mannosides. J. Med. Chem. 55, 3945–3959 (2012).

29. Spaulding, C. N. et al. Selective depletion of uropathogenic E. coli from the gut by a FimH antagonist. Nature 546, 528–532 (2017).

30. Hartmann, M. & Lindhorst, T. K. The Bacterial Lectin FimH, a Target for Drug Discovery - Carbohydrate Inhibitors of Type 1 Fimbriae-Mediated Bacterial Adhesion. Eur. J. Org. Chem. 2011, 3583–3609 (2011).

31. Appeldoorn, C. C. M. et al. Novel multivalent mannose compounds and their inhibition of the adhesion of type 1 fimbriated uropathogenic E. coli. Tetrahedron Asymmetry 16, 361–372 (2005).

32. Jarvis, C. et al. Antivirulence Isoquinolone Mannosides: Optimization of the Biaryl Aglycone for FimH Lectin Binding Affinity and Efficacy in the Treatment of Chronic UTI. ChemMedChem 11, 367–373 (2016).

33. Mydock-McGrane, L. et al. Antivirulence C-Mannosides as Antibiotic-Sparing, Oral Therapeutics for Urinary Tract Infections. J. Med. Chem. 59, 9390–9408 (2016).

34. Nilsson, L. M., Thomas, W. E., Trintchina, E., Vogel, V. & Sokurenko, E. V. Catch Bond-mediated Adhesion without a Shear Threshold: Trimannose Versus Monomannose Interactions With The FimH Adhesin of Escerichia coli. J. Biol. Chem. 281, 16656–16663 (2006).

35. Björnham, O., Nilsson, H., Andersson, M. & Schedin, S. Physical properties of the specific PapG–galabiose binding in E. coli P pili-mediated adhesion. Eur. Biophys. J. 38, 245 (2009).

36. Duncan, M. J. et al. The Distinct Binding Specificities Exhibited by Enterobacterial Type 1 Fimbriae Are Determined by Their Fimbrial Shafts. J. Biol. Chem. 280, 37707–37716 (2005).

37. Spaulding, C. N. et al. Functional role of the type 1 pilus rod structure in mediating host-pathogen interactions. eLife 7, e31662 (2018).

38. Reisner, A. et al. Type 1 Fimbriae Contribute to Catheter-Associated Urinary Tract Infections Caused by Escherichia coli. J. Bacteriol. 196, 931–939 (2014).

39. Rodrigues, D. F. & Elimelech, M. Role of type 1 fimbriae and mannose in the development of Escherichia coli K12 biofilm: from initial cell adhesion to biofilm formation. Biofouling 25, 401–411 (2009).

40. Wang, L. et al. Influence of Type I Fimbriae and Fluid Shear Stress on Bacterial Behavior and Multicellular Architecture of Early Escherichia coli Biofilms at Single-Cell Resolution. Appl. Environ. Microbiol. 84, e02343–17 (2018).

41. Liang, M. N. et al. Measuring the forces involved in polyvalent adhesion of uropathogenic Escherichia coli to mannose-presenting surfaces. Proc. Natl. Acad. Sci. 97, 13092–13096 (2000).

42. Curk, T., Dobnikar, J. & Frenkel, D. Design Principles for Super Selectivity using Multivalent Interactions. in Multivalency 75–101 (John Wiley & Sons, Ltd, 2017). doi:10.1002/9781119143505.ch3.

43. Björnham, O. & Axner, O. Catch-Bond Behavior of Bacteria Binding by Slip Bonds. Biophys. J. 99, 1331–1341 (2010).

44. Whitfield, M., Ghose, T. & Thomas, W. Shear-Stabilized Rolling Behavior of E. coli Examined with Simulations. Biophys. J. 99, 2470–2478 (2010).

45. Hammer, D. A. & Apte, S. M. Simulation of cell rolling and adhesion on surfaces in shear flow: general results and analysis of selectin-mediated neutrophil adhesion. Biophys. J. 63, 35–57 (1992).

46. Dasanna, A. K., Lansche, C., Lanzer, M. & Schwarz, U. S. Rolling Adhesion of Schizont Stage Malaria-Infected Red Blood Cells in Shear Flow. Biophys. J. 112, 1908–1919 (2017).

47. Ye, H., Shen, Z. & Li, Y. Cell Stiffness Governs Its Adhesion Dynamics on Substrate Under Shear Flow. IEEE Trans. Nanotechnol. 17, 407–411 (2018).

48. Wu, T.-H. & Qi, D. Investigation of shear rates of rolling adhesion on leukocytes with bending of microvilli. Phys. Rev. Fluids 4, 063101 (2019).

49. Luo, Z. Y. & Bai, B. F. State diagram for adhesion dynamics of deformable capsules under shear flow. Soft Matter 12, 6918–6925 (2016).

50. Farokhirad, S. et al. Stiffness can mediate balance between hydrodynamic forces and avidity to impact the targeting of flexible polymeric nanoparticles in flow. Nanoscale 11, 6916–6928 (2019).

51. Davies, H. S. et al. An integrated assay to probe endothelial glycocalyx-blood cell interactions under flow in mechanically and biochemically well-defined environments. Matrix Biol. 78–79, 47–59 (2019).

52. Molaei, M., Barry, M., Stocker, R. & Sheng, J. Failed Escape: Solid Surfaces Prevent Tumbling of *Escherichia coli*. Phys. Rev. Lett. 113, 068103 (2014).

53. Cheong, F. C. et al. Rapid, High-Throughput Tracking of Bacterial Motility in 3D via Phase-Contrast Holographic Video Microscopy. Biophys. J. 108, 1248–1256 (2015).

54. Molaei, M. & Sheng, J. Succeed escape: Flow shear promotes tumbling of Escherichia colinear a solid surface. Sci. Rep. 6, 35290 (2016).

55. Bianchi, S., Saglimbeni, F. & Leonardo, R. D. Holographic Imaging Reveals the Mechanism of Wall Entrapment in Swimming Bacteria. Phys. Rev. X 7, 011010 (2017).

56. Öberg, K. et al. Templating Gold Surfaces with Function: A Self-Assembled Dendritic Monolayer Methodology Based on Monodisperse Polyester Scaffolds. Langmuir 29, 456–465 (2013).

57. McMichael, J. C. & Ou, J. T. Structure of common pili from Escherichia coli. J. Bacteriol. 138, 969–975 (1979).

58. Lundgren, A., et al. Self-Assembled Arrays of Dendrimer-Gold-Nanoparticle Hybrids for Functional Cell Studies. Angew. Chem. Int. Ed. 50, 3450–3453 (2011).

59. Bell, G. Models for the specific adhesion of cells to cells. Science 200, 618–627 (1978).

60. Le Trong, I. et al. Structural Basis for Mechanical Force Regulation of the Adhesin FimH via Finger Trap-like β Sheet Twisting. Cell 141, 645–655 (2010).

61. Loferer-Krößbacher, M., Klima, J. & Psenner, R. Determination of Bacterial Cell Dry Mass by Transmission Electron Microscopy and Densitometric Image Analysis. Appl. Environ. Microbiol. 64, 688–694 (1998).

62. Sjollema, J. et al. Detachment and successive re-attachment of multiple, reversibly-binding tethers result in irreversible bacterial adhesion to surfaces. Sci. Rep. 7, 4369 (2017).

63. Vissers, T. et al. Bacteria as living patchy colloids: Phenotypic heterogeneity in surface adhesion. Sci. Adv. 4, eaao1170 (2018).

64. Schwartz, D. J. et al. Positively selected FimH residues enhance virulence during urinary tract infection by altering FimH conformation. Proc. Natl. Acad. Sci. 110, 15530–15537 (2013).

65. Anderson, B. N. et al. Weak Rolling Adhesion Enhances Bacterial Surface Colonization. J. Bacteriol. 189, 1794–1802 (2007).

66. Schluter, J., Nadell, C. D., Bassler, B. L. & Foster, K. R. Adhesion as a weapon in microbial competition. ISME J. 9, 139–149 (2015).

67. Zhao, K. et al. Psl trails guide exploration and microcolony formation in Pseudomonas aeruginosa biofilms. Nature 497, 388–391 (2013).

68. Nadell, C. D., Ricaurte, D., Yan, J., Drescher, K. & Bassler, B. L. Flow environment and matrix structure interact to determine spatial competition in Pseudomonas aeruginosa biofilms. eLife 6, e21855 (2017).

69. Armbruster, C. R. et al. Heterogeneity in surface sensing suggests a division of labor in Pseudomonas aeruginosa populations. eLife https://elifesciences.org/articles/45084 (2019) doi:10.7554/eLife.45084.

70. Nadell, C. D. & Bassler, B. L. A fitness trade-off between local competition and dispersal in Vibrio cholerae biofilms. Proc. Natl. Acad. Sci. 108, 14181–14185 (2011).

71. Schembri, M. A., Pallesen, L., Connell, H., Hasty, D. L. & Klemm, P. Linker insertion analysis of the FimH adhesin of type 1 fimbriae in an Escherichia coli fimH-null background. FEMS Microbiol. Lett. 137, 257–263 (1996).

72. Klemm, P., Jrgensen, B. J., van Die, I. & de Ree, H. The Fim genes responsible for synthesis of type 1 fimbriae in Escherichia coli, cloning and genetic organization. Mol. Genet. Genomics 199, 410–414 (1985).

73. Schembri, M. A., Christiansen, G. & Klemm, P. FimH-mediated autoaggregation of Escherichia coli. Mol. Microbiol. 41, 1419–1430 (2001).

74. Schneider, C. A., Rasband, W. S. & Eliceiri, K. W. NIH Image to ImageJ: 25 years of image analysis. Nat. Methods 9, 671–675 (2012).

75. Lundgren, A. et al. Gold-Nanoparticle-Assisted Self-Assembly of Chemical Gradients with Tunable Sub-50 nm Molecular Domains. Part. Part. Syst. Charact. 31, 209–218 (2014).

76. Slot, J. W. & Gueze, H. J. Sizing of protein A-colloidal gold probes for immunoelectron microscopy. J. Cell Biol. 90, 533 (1981).

77. Hulander, M., Valen-Rukke, H., Sundell, G. & Andersson, M. Influence of Fibrinogen on *Staphylococcus epidermidis* Adhesion Can Be Reversed by Tuning Surface Nanotopography. ACS Biomater. Sci. Eng. 5, 4323–4330 (2019).

78. Lima, R., Wada, S., Tsubota, K. & Yamaguchi, T. Confocal micro-PIV measurements of three-dimensional profiles of cell suspension flow in a square microchannel. Meas. Sci. Technol. 17, 797–808 (2006).

79. van Oostrum, P. D. J. & Reimhult, E. O. A Method for Determining a Three-Dimensional Particle Distribution in a Medium. (2019).

80. Bruus, H. Governing Equations in Microfluidics. in Microscale Acoustofluidics 1–28 (Royal Society of Chemistry, 2014).

81. Madsen, K., Nielsen, H. B. & Tingleff, O. Methods for Non-Linear Least Squares Problems (2nd Ed.). https://orbit.dtu.dk/en/publications/methods-for-non-linear-least-squares-problems-2nd-ed (2004).

