## Supplemental information for "Adhesion of *E. coli* bacteria is force-modulated due to fimbriae-mediated surface repulsion and multivalent binding irrespective of surface specificity"

**List of contents**

Supplementary figure 1:
Samples with immobilized *E. coli* used for AFM experiments p.2

Supplementary figure 2:
Full approach-and-retract force spectra obtained with AFM p.3

Supplementary figure 3:
Surface concentration of gold nanoparticles obtained for different adsorption conditions visualized by electron microscopy p.4

Supplementary figure 4:
Calculated flow profiles in the microfluidic channels used for bacterial binding experiments recorded with brightfield microscopy p.5

Supplementary figure 5:
Distribution of binding times measured for *E. coli* binding to different nanopatterns determined from footage acquired with brightfield microscopy p.6

Supplementary calculation:
Estimation of forces acting on surface tethered bacteria subject to flow p. 7

**Supplementary Figure 1: Samples with immobilized *E. coli* used for AFM experiments**

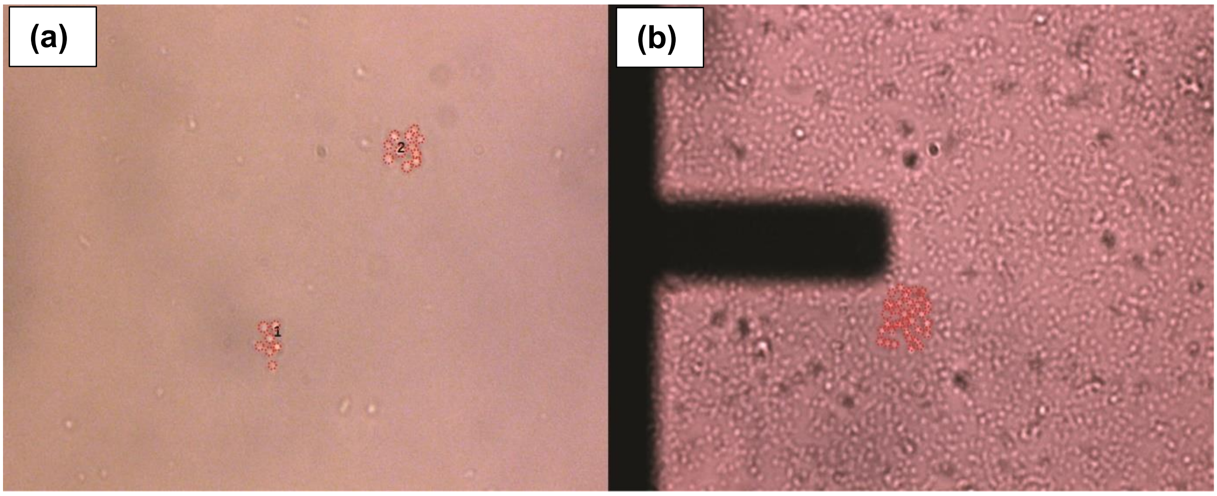


The figure shows two images that outline groups of immobilized *E. coli* bacteria that were probed with AFM using **(a)** mannosylated cantilevers and **(b)** with hydrophobic cantilevers as detailed in the method section. Individual bacteria within each group are indicated by red circles. For the case of high surface coverage of bacteria, the number of encircled bacteria is an approximation based on the contact area of the cantilever. The right image also shows the contours of the tip of the cantilever, slightly out of focus. The images were obtained just before AFM analysis with the optical microscope (Axio Observer Z1, Zeiss, Germany) onto which the AFM measurement module (JPK Nanowizard III, Bruker Nano GmbH, Germany) had been mounted.

**Supplementary Figure 2: Full approach-and-retract force spectra obtained with AFM**

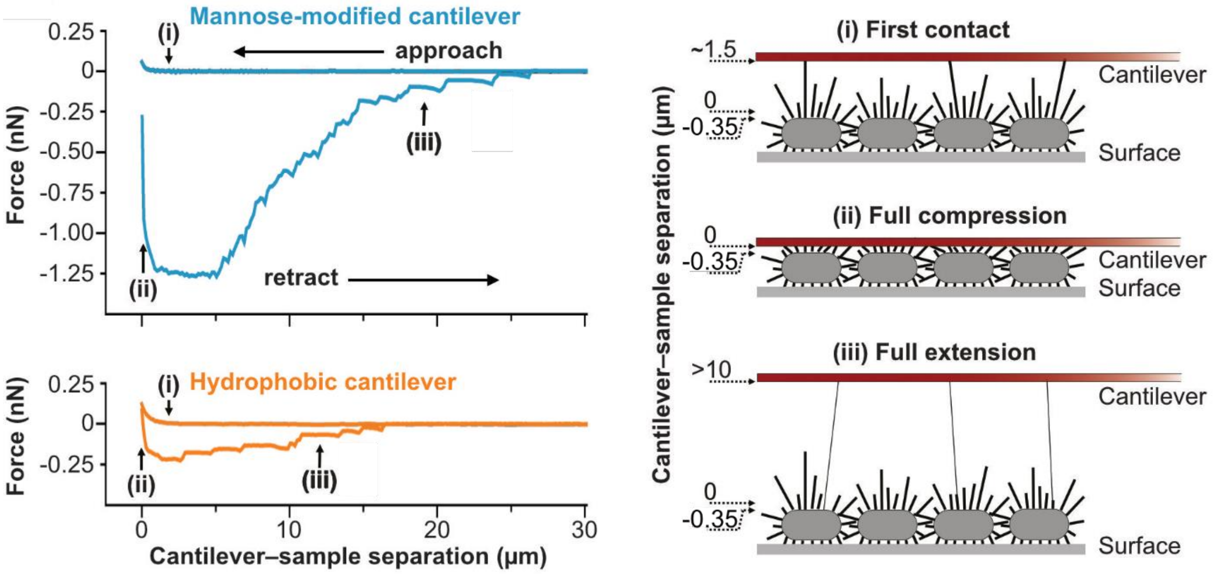


The graphs to the left show two examples of full force spectra obtained with a mannosylated cantilever (upper spectrum, blue line) and a hydrophobic cantilever (lower spectrum, orange line) when first approaching a group of surface-immobilized *E. coli* bacteria, halting the cantilever 0.35 µm above the bacterial surfaces for <0.5 s. (note that this position corresponds to cantilevers-sample separation zero in the spectra) and then retracting the cantilever until all attached fimbriae had released. The cartoon to the right explains our interpretation of the spectra at three different points during the measurement.

**Supplementary Figure 3: Surface concentration of gold nanoparticles obtained for different adsorption conditions visualized by electron microscopy**

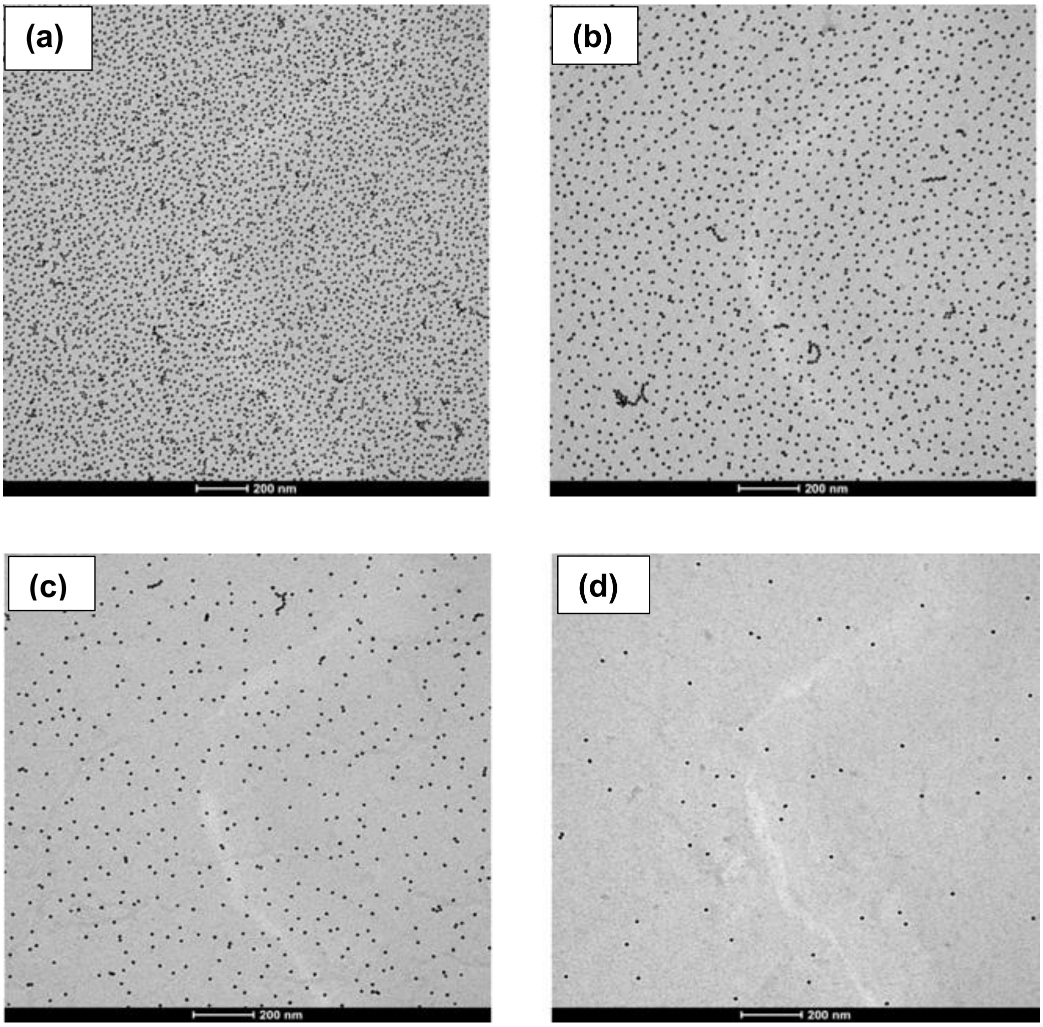


The figure shows four Transmission Electron Micrographs (TEM) displaying the distribution and surface concentration of 10-nm gold nanoparticles (NPs) adsorbed under different conditions to silicon dioxide coated TEM grids (SF300-CU, Electron Microscopy Sciences) treated with (3-aminopropyl)dimethylethoxysilane (APDMES 95%, ABCR, Germany). The grids were first cleaned/pre-treated in a UV/ozone chamber (ProCleaner, Bioforce Nanoscience, USA) for 15 minutes and then modified with APDMES from gaseous phase. This was done by placing the grids in a closed container together with a small amount of APDMES solution (50% in Methanol) for 30 minutes whereupon the grids were washed gently first with methanol, then with water, and finally dried under a gentle stream of $N_2$. Gold nanoparticles, ~10 nm in diameter (synthesized as detailed in the method section), were bound to the grids using the same conditions as was used to modify the microfluidic channels, i.e., by immersing grids for 15 minutes in solution containing **(A)** 30 nM NPs in 10 mM citric buffer pH4.0 (CB), **(B)** 30 nM NPs in 0.08 mM CB, **(C)** 3 nM NPs in 0.08 mM CB, and **(D)** 0.3 nM NPs in 0.08 mM CB. Electron micrographs were captured using a FEI Tecnai G2 microscope operated at 160 kV acceleration voltage.

**Supplementary Figure 4: Calculated flow profiles in the microfluidic channels used for bacterial binding experiments recorded with brightfield microscopy**

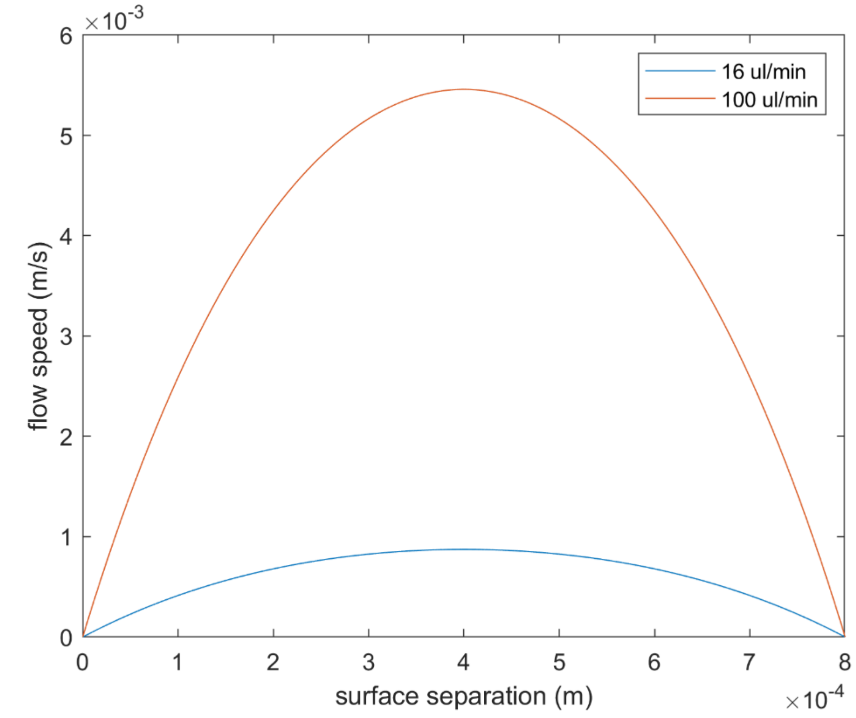


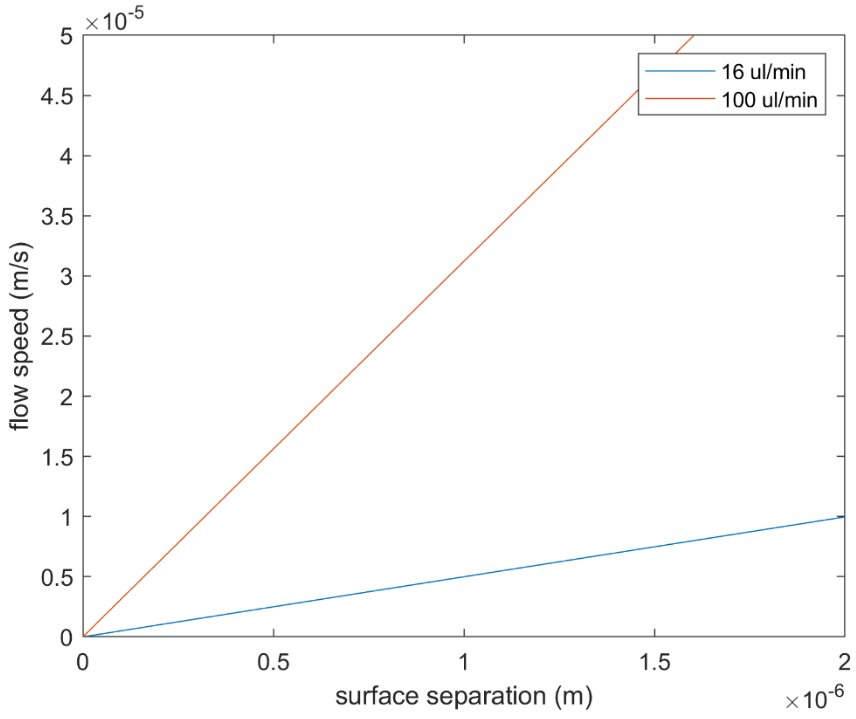


The graphs show the flow speed at different separations from the bottom of a square microfluidic channel with width (W) and height (H) 800 µm. The flow profiles were calculated according to Lima *et al.* (*Meas. Sci. Technol.* 2006, **17**, 797–808) for volumetric flows of 16 µl min$^{-1}$ and 100 µl min$^{-1}$ at the center of the channel. Plot **(a)** shows the full flow profiles and plot (**b**) highlights the flow profiles close to the channel floor.

**Supplementary figure 5: Distribution of binding times measured for *E. coli* binding to different nanopatterns determined from footage acquired with brightfield microscopy**

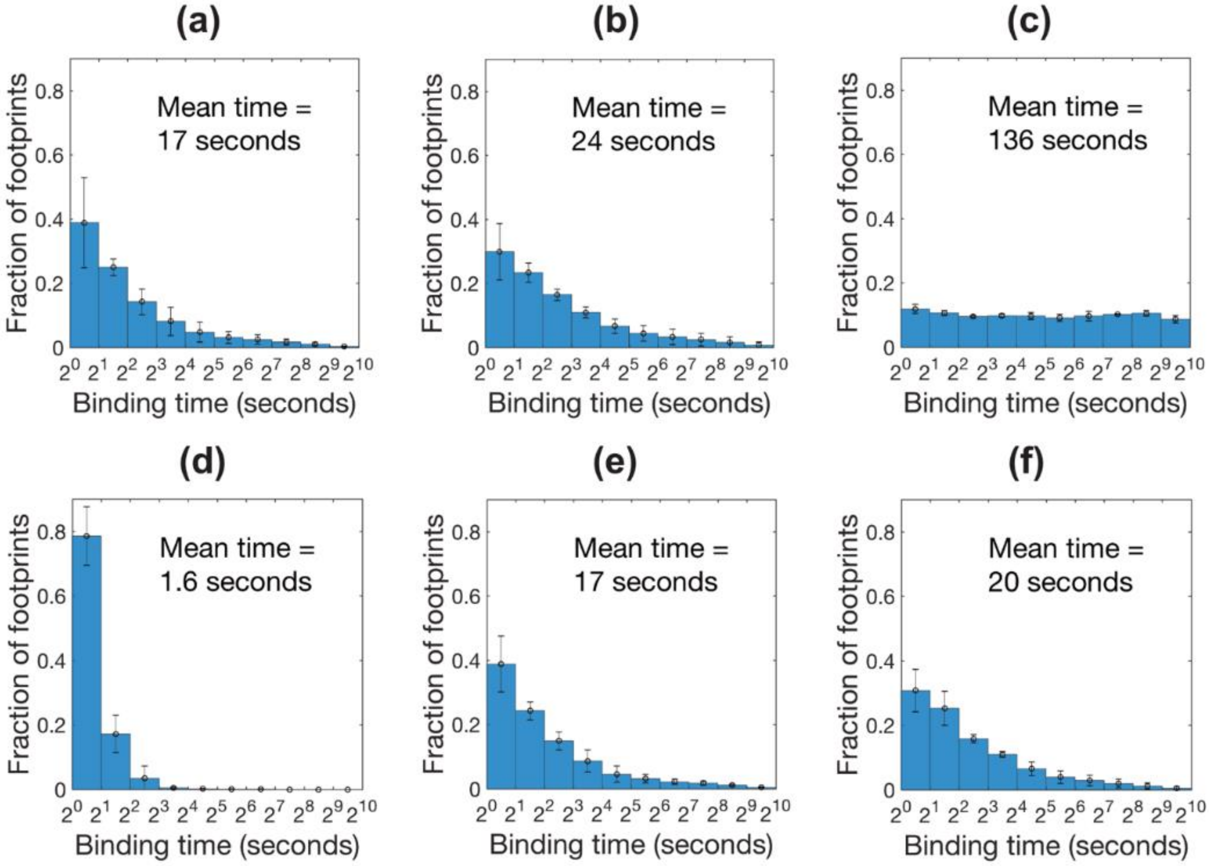


The graphs show the distribution of binding times detected for binding of fimbriated *E. coli* to nanopatterns displaying (a) <10 mannose-modified NPs µm$^{-2}$, (b) 30 mannose-modified NPs µm$^{-2}$, 150 mannose-modified NPs µm$^{-2}$, (d), 150 hydrophobic NPs µm$^{-2}$, 650 hydrophobic NPs µm$^{-2}$ and 1500 hydrophobic NPs µm$^{-2}$. Data were acquired using a volumetric flow of 16 µl min$^{-1}$ corresponding to a flow speed of 5 µm s$^{-s}$ at 1 µm surface separation. Note that a logarithmic scale with base 2 is used for the time axis. Each bar indicates the average and standard deviation of ≥3 independent experiments where each experiment includes approximately 1000 footprints.

**Supplementary calculation: Estimation of forces acting on surface tethered bacteria subject to flow**

A surface-tethered bacterium exposed to liquid flow will be subject to a hydrodynamic drag force. According to Stokes' law this force is given by $\mathrm{F_F} = \gamma \times \mathrm{v}$ where v is the flow speed relative to the speed of the bacterium and $\gamma$ is the friction factor given by $\gamma = 6\pi\eta\mathrm{R}$ where $\eta$ is the dynamic viscosity of the liquid and R is the effective radius of the bacterium. To a first approximation we assume that $\mathrm{R} \approx 1\ \mu\mathrm{m}$, giving $\gamma = 1.9 \times 10^{-8}\ \mathrm{kg\ s^{-1}}$, and that $\mathrm{v} \approx \mathrm{v}_z$, the local flow speed at the separation z from the surface plane of the microfluidic channel. The hydrodynamic drag force acting on a bacterium bound at separation z can thus be estimated by $\mathrm{F_F} = 1.9 \times 10^{-8} \times \mathrm{v}_z$. For tethered bacteria $\mathrm{z} < 2\ \mu\mathrm{m}$ and therefore $\mathrm{v_z} < 60\ \mu\mathrm{m\ s^{-1}}$ when the volumetric flow is $100\ \mu\mathrm{L\ min^{-1}}$ (Supplementary figure 4). Accordingly, we can assume that $\mathrm{F_F} < 1.2\ \mathrm{pN}$ for the conditions used in the bacterial binding experiments.

Because of the drag force $\mathrm{F_F}$ acting on a surface-tethered bacterium there will be a torque and, by the combination of the tensile force in the fimbria, a force normal to the surface plane $\mathrm{F_N}$ that pushes the bacterium towards the surface. Under the assumption that the tether initially is oriented approximately normal to the surface, the magnitude of $\mathrm{F_N}$ associated with the displacement towards the surface can be estimated from knowledge of the positions of the bacterium before $(\mathrm{Z_1})$ and after $(\mathrm{Z_2})$ and the hydrodynamic drag force acting on it when positioned at $\mathrm{z} = \mathrm{Z_2}$. First one calculates the force perpendicular to the tether as $\mathrm{F_T} = \mathrm{F}_F \sin\theta = \mathrm{F}_F \times (\mathrm{Z_2}/\mathrm{Z_1})$ of which the component normal to the surface is $\mathrm{F_N} = \mathrm{F_T}\cos\theta$, as detailed in the scheme below. DHPCM data showed that the bacteria relocate from $\mathrm{Z_1} = 1.6\ \mu\mathrm{m}$ to $\mathrm{Z_2} = 1.0\ \mu\mathrm{m}$ when the flow speed increases to $\mathrm{v_{Z_2}} = 35\ \mu\mathrm{m\ s^{-1}}$. Using these values in the above model yields $\mathrm{F_N} = 0.32\ \mathrm{pN}$.

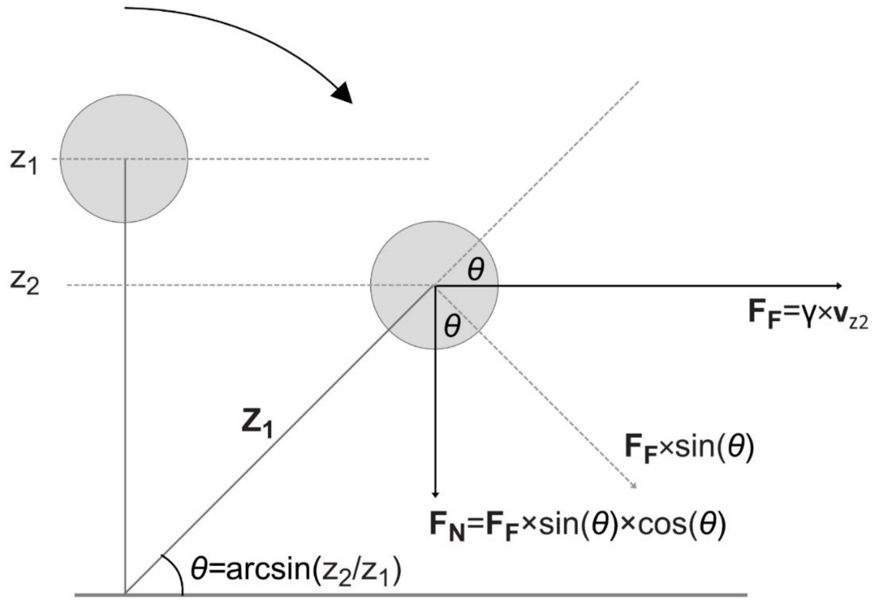
